# Taste receptor T1R3 in nasal cilia detects *Staphylococcus aureus* D-amino acids to increase apical glucose uptake and enhance innate immunity

**DOI:** 10.1101/2022.10.21.512955

**Authors:** Ryan M. Carey, Benjamin M. Hariri, Nithin D. Adappa, James N. Palmer, Robert F. Margolskee, Noam A. Cohen, Robert J. Lee

## Abstract

Bitter (T2R) and sweet/umami (T1R) taste receptors serve chemosensory roles throughout the body. In airway cilia, T2Rs detect bacterial metabolites to stimulate bactericidal nitric oxide. T1Rs in solitary chemosensory cells detect glucose in airway surface liquid (ASL) and bacterial D-stereoisomer amino acids to regulate antimicrobial peptides. Using differentiated air-liquid interface cultures of primary nasal cells, we show that the T1R3 receptor is also expressed in human and mouse nasal cell cilia. D-amino acids produced at low mM concentrations by *Staphylococcus aureus* activate T1R3 to decrease ASL glucose and increase apical glucose uptake by increasing GLUT2 and GLUT10 expression through a β-arrestin pathway. Our data suggests that T1R3 localized to cilia functions as an immune detector for D-amino acids to reduce ASL glucose and potentially limit bacterial growth. Given that glucose protected *S. aureus* against bactericidal NO produced during T2R activation, the reduced ASL glucose with T1R3 activation may also sensitize bacteria to other innate defenses.

**HIGHLIGHTS:**

- Nasal motile cilia express the T1R3 subunit of the sweet taste receptor
- *S. aureus* D-amino acids activate cilia T1R3 to enhance mucosal glucose uptake
- Reduced airway surface liquid glucose likely reduces *S. aureus* growth
- Reduced airway glucose also sensitizes *S. aureus* to epithelial NO production

## INTRODUCTION

The regulation of airway surface liquid (ASL) glucose is an important component of epithelial innate defense. Despite constant diffusional glucose flux across the airway epithelial barrier from the serosal fluid into the airway lumen,^1–3^ ASL glucose is kept ∼10-fold lower than serum glucose to limit glucose availability for bacterial growth.^4–6^ Airway cells keep the ASL glucose low through apical re-uptake of airway glucose through glucose transporters, including GLUT2 and/or GLUT10,^7,8^ followed by epithelial metabolism of the glucose.^9^ Hyperglycemia can increase the driving force for glucose flux, and inflammation can disrupt tight junctions to increase glucose leak.^7,10^ Both conditions can increase ASL glucose and create a more favorable environment for pathogen growth.^1,2,4,11,12^ We found that patients with chronic rhinosinusitis (CRS) have increased glucose in their nasal secretions compared with control individuals, despite no differences in blood glucose levels.^13^ Elevated ASL glucose in CRS patients is not lowered by the use of corticosteroids,^14^ and thus reducing excess nasal fluid glucose may be a beneficial complementary therapeutic strategy regardless of a patient’s responsiveness to steroid therapy.

We previously showed that elevated ASL glucose can inhibit innate defense responses mediated by solitary chemosensory cells (SCCs). SCCs are also known as “tuft cells” because of similarities to intestinal tuft cells named for their apical tuft of microvilli. These cells are also referred to as “brush cells.”^15^ SCCs regulate several responses, including IL-25 and/or acetylcholine release to activate epithelial inflammation or trigger mucus clearance, respectively.^16,17^ In the nose, they also regulate secretion of antimicrobial peptides like defensins.^13,18^ SCC activation of antimicrobial peptide secretion is reduced by activation of the sweet taste receptor (taste family 1 receptor) subunits T1R2 and T1R3 via ASL glucose^13^ or sweet bacterial D-amino acids.^18,19^ While eukaryotes have evolved to use primarily L-stereoisomer amino acids, many bacteria produce and use D-amino acid stereoisomers.^20,21^ Many D-amino acids activate the sweet taste receptor and thus taste sweet.^22^ Both D-Phe and D-Ile produced by *Staphylococcus aureus* can activate SCC sweet taste receptors and inhibit antimicrobial peptide release in primary nasal cells *in vitro*.^18^

Airway motile cilia are critical to clear the respiratory tract of inhaled particulates and pathogens that get trapped in airway mucus within the ASL.^23^ Coordinated beating of cilia transports mucus to the oropharynx, where it is expectorated or swallowed. More recently, airway motile cilia have also been identified as chemosensory organelles important in cell signaling.^24–26^ In nasal epithelial cells, bitter taste receptors (taste family 2 receptors or T2Rs) in airway cilia detect bitter bacterial lactones and quinolones to activate nitric oxide (NO) production to clear and kill bacteria.^27–29^

A previous study of rat tracheal epithelium suggested that the T1R3 sweet taste receptor subunit is expressed in multi-ciliated cells as well as SCCs.^30^ This study demonstrated that T1R3 was co-expressed in cells with GLUT2, and thus the authors hypothesized that T1R3 in ciliated cells may play a role in glucose transport. Given the role of ASL glucose regulation in immune defense, we hypothesized that T1R3 in airway cilia may function in part to sense “sweet” bacterial metabolites like D-amino acids to increase glucose uptake and reduce pathogen growth. Here, we sought to determine if human nasal epithelial cells express the T1R3 sweet taste receptor subunit within motile cilia, and if this receptor regulates glucose transport in response to bacterial D-amino acids. The sweet taste receptor is highly druggable via artificial sweetener agonists^31,32^ or antagonists like lactisole^33^ and gymnemic acid,^34^ many of which are already approved for ingestion and could be delivered via topical nasal rinse.^35^ We thus also hypothesize that the sweet taste receptor may be a logistically feasible target for altering nasal epithelial glucose concentrations in some CRS patients.

## RESULTS

### T1R3 expression and localization to motile cilia and negative regulation by TLR5

T1R3 expression was detected by immunofluorescence in human motile cilia co-labeled with β-tubulin IV primary antibody (**Figure 1A**) using rabbit serum as a non-specific binding control (**Figure 1B**). While we previously published evidence of T1R3 expression in SCCs,^13,18^ the cilia-localized T1R3 was not initially apparent as cilia are more difficult to permeabilize. For IF of SCCs, we typically utilize gentler permeabilization (0.1% triton, 10 min) than for cilia (0.3% triton, 1 hour), as cilia are covered with cell attached mucins which make permeabilization difficult. Orthogonal slices confirmed expression of T1R3 at the level of cilia (βTubIV) above the level of phalloidin, which stains the actin-enriched microvilli at the cilia base (**Figure 1C**). Direct primary antibody labelling showed overlapping staining of two different T1R3 antibodies (**Figure 1D**). Cilia staining with a third T1R3 antibody was reduced by its corresponding antigenic blocking peptide (**Figure 1E**). Similar staining was observed in whole mount fresh nasal tissue (**Figure 1F**). Finally, we observed cilia staining with an anti-mouse T1R3 antibody in mouse nasal septal ALIs, and this staining was absent in ALIs derived from tissue from T1R3 knockout (T1R3^-/-^) mice (**Figure 1G**).

**Figure 1.**
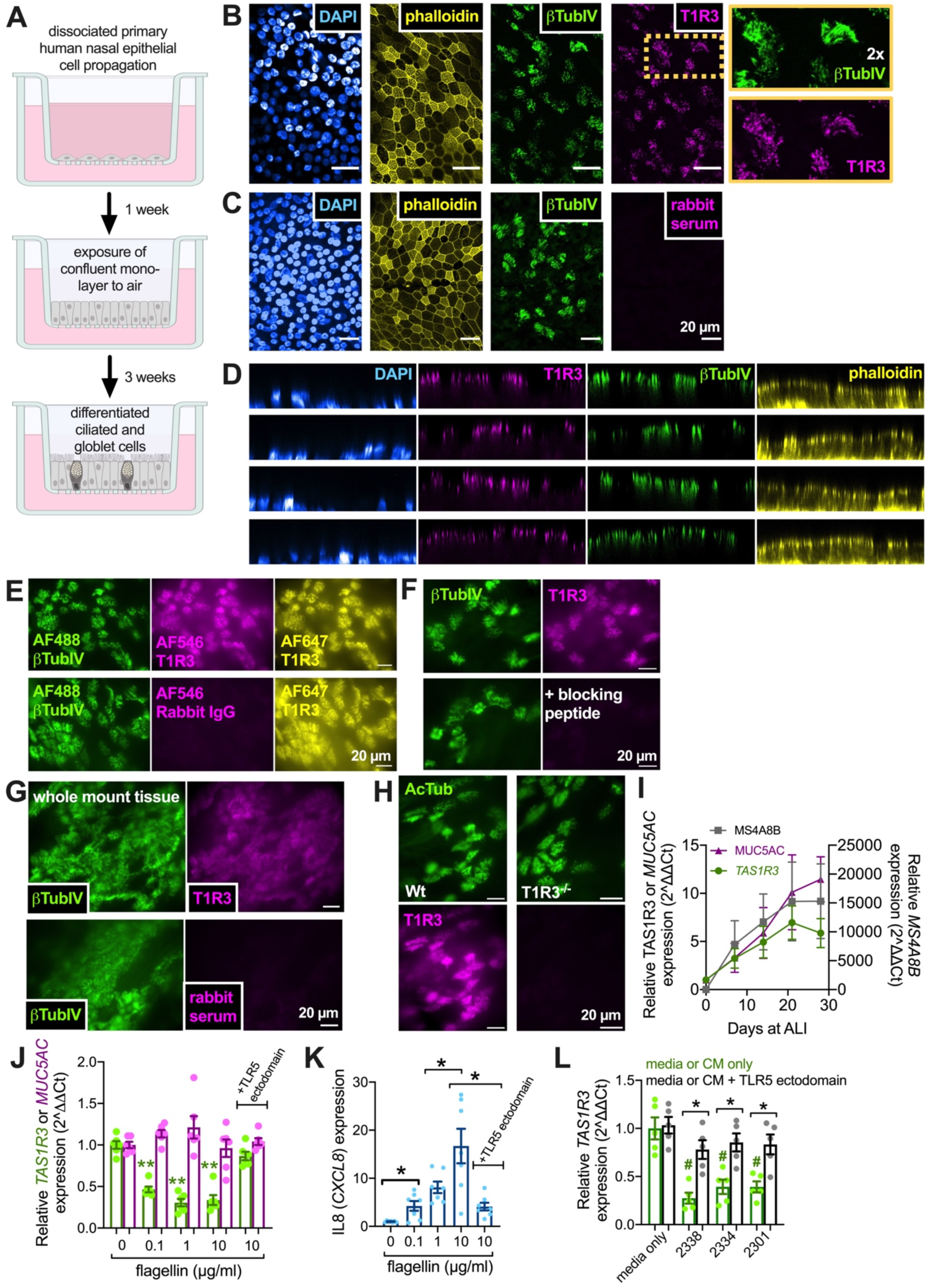
T1R3 localization to nasal epithelial cilia and *TAS1R3* mRNA expression in nasal ALI cultures. Primary nasal tissue is enzymatically dissociated and basal epithelial cells are propagated in submersion culture. Cells on permeable Transwell filters are grown to confluence and the medium on the apical surface is removed. The resulting air-liquid interface (ALI) culture leads to the differentiation of ciliated and goblet cells mimicking the *in vivo* epithelium. Image created in Biorender. **(B)** Laser-scanning confocal micrographs of immunofluorescence of T1R3 (magenta) and β-tubulin IV (pink) taken at the same plane in nasal ALI culture. Phalloidin (taken at a plane just below the cilia; yellow) and DAPI (nuclear stain; blue; taken at a plane lower than the phalloidin) shown as a control for epithelial integrity. Boxes on the right show the region outlined in orange at 2x to show co-localization of T1R3 and β-tubulin IV. **(C)** Control staining with rabbit serum showed no cilia labeling. Results in A-B representative of staining observed in n = 3 ALIs from individual patients. **(D)** Orthogonal slices (*xz*) from immunostaining experiments as in *A* showing the co-localization of T1R3 (magenta) and β-tubulin IV (green) above the level of the phalloidin (yellow). Antibody used in *A* and *C* was Abcam ab150525. **(E)** Direct labeling with AlexaFluor Zenon labeling kits allowed co-staining with AF546-labeled anti-T1R3 (PA5-34256; ThermoFisher; magenta) and AF647-labeled anti-T1R3 (SAB4503300; Sigma; yellow), which both showed co-labeling of cilia (β-tubulin IV; green). Labeling of Rabbit IgG control with AF546 showed no fluorescence of cilia. Results representative of staining observed in n = 3 ALIs from individual patients. **(F)** Cilia staining with another T1R3 antibody (MBS421865; MyBioSource) was reduced by its antigenic blocking peptide (10-fold molar excess; 1 hour pre-incubation). Results representative of staining observed in n = 3 ALIs from individual patients. **(G)** We also saw cilia-staining of T1R3 (ab150525) in thin mucosal tissue pieces acutely removed during sinus surgery. Results representative of staining observed in tissue from n = 3 from individual patients. **(H)** A mouse-directed T1R3 antibody (NB100-98792) stained cilia in ALIs derived from Wt mouse nasal septum but not ALIs derived from *TAS1R3^-/-^* mouse septum. All immunofluorescence results representative of staining observed in n = 3 from ALIs from each genotype. **(I)** Graph showing expression of *TAS1R3* (green), goblet cell marker *MUC5AC* (magenta), and cilia marker *MS4A8B* by qPCR at time points after exposure of ALI cultures to air. **(J-K)** Bar graphs showing *TAS1R3* and *MUC5AC* (*B*) or *IL8* (*CXCL8; C*) expression at day 21 ALIs ± TLR5 agonist flagellin ± soluble TLR5 ectodomain antagonist. **(K)** Graph showing *TAS1R3* expression after exposure to conditioned media (CM) from patients with *P. aeruginosa* respiratory infections ± TLR5 ectodomain. All data are presented as mean ± SEM using ALIs from n = 6 (*A* and *B*), 7 (*C*), or 5 (*D*) patients in independent experiments. Significances in *B*-*D* determined by one way ANOVA with Bonferroni posttest; \**p*<0.05.

We saw an increase in *TAS1R3* gene transcript by qPCR as primary nasal cells were differentiated at ALI (**Figure 1H**), as previously observed with cilia-expressed *TAS2R* bitter receptor genes.^29^ Goblet-cell *MUC5AC* and cilia-localized *MS4A8B*^36^ were used as controls. As we also previously observed with several *TAS2R*s,^29^ *TAS1R3* transcript was reduced by airway cell stimulation with toll-like receptor 5 (TLR5) agonist *Pseudomonas aeruginosa* flagellin (**Figure 1I**). Both *TAS1R3* decrease and CXCL8 (IL-8) increase were reduced by addition of soluble TLR5 ectodomain (**Figure 1I-J**), confirming this was due to flagellin activation of TLR5. Conditioned media from bacteria isolated from three CRS patients with confirmed *Pseudomonas aeruginosa* infections likewise decreased *TAS1R3* expression in primary nasal ALIs (**Figure 1K**). These data suggest that T1R3 is expressed and localized to nasal cilia and is negatively regulated by TLR5.^29^

### T1R3 modulates airway surface liquid (ASL) glucose

ASL glucose regulation is a balance of glucose leak and glucose re-uptake by epithelial cells (**Figure 2A**). When cultured at ALI, cell lines like RPMI2650 or A549 cancer cells form less of a tight junction barrier (evidenced by lower transepithelial electrical resistance [TEER]; **Figure 2B**) compared with primary nasal cells and are less able to keep ASL glucose concentrations low during acute (18 hr) challenge of high glucose (**Figure 2C**). We noted that, in the presence of high basolateral glucose (10 or 15 mM), primary nasal cells also exhibit increased ASL glucose in the presence of T1R3 allosteric inhibitor lactisole (2 mM; apical side only; 18 hrs.) ^32,33^ and lower ASL glucose in the presence of non-metabolizable T1R3 agonist sucralose (2 mM; apical side only; 18 hrs.) (**Figure 2D**). There was no change in TEER (**Figure S1A**), FITC-dextran permeability (**Figure S1B**), or IL-8 transcription (**Figure S1C**) with sucralose ± lactisole treatment. This suggests sucralose and lactisole cause neither a reduction in barrier function, which would be reflected by reduced TEER, nor an increase in inflammatory state of the cells, which would be reflected in NFκB-driven IL-8 production.

**Figure 2.**
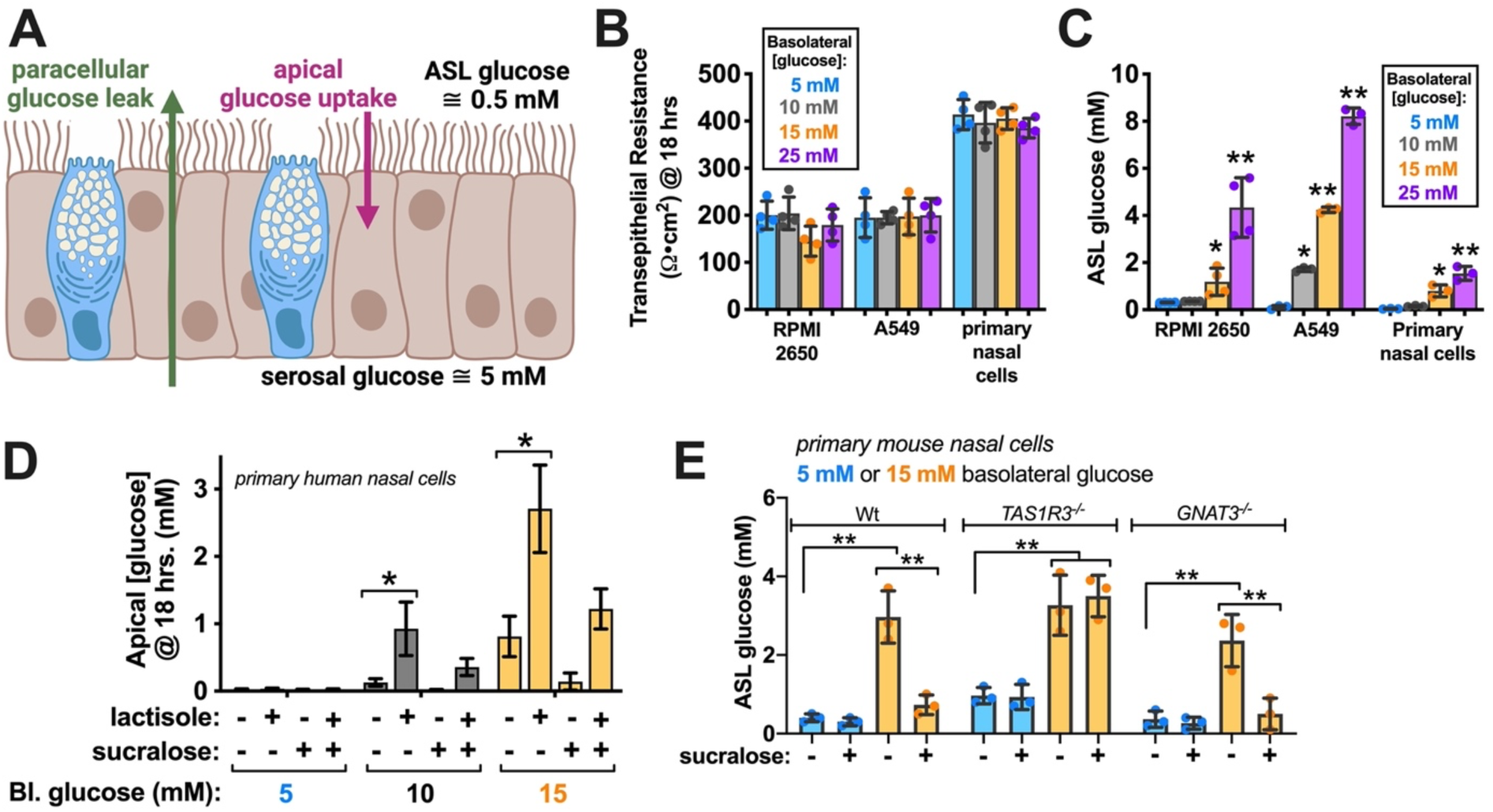
T1R3 regulates airway surface liquid (ASL) glucose. **(A)** Model of ASL glucose concentration as a balance between tonic leak from serosal fluid and apical uptake via GLUT transporters from ref^2^. Goblet cells in blue and ciliated cells in brown. In healthy airways, ASL glucose is kept ∼10-fold lower than serosa. Image created with Biorender. **(B-C)** Bar graph showing transepithelial resistance (TEER; *B*) and apical glucose (*C*) of ALI cultures grown from cell lines (RPMI2650, A549) or primary nasal epithelial cells after incubation in 5, 10, 15, and 25 mM glucose; n = 4 independent experiments per condition, primary cells utilized ALIs from 4 individual patients. **(D)** Bar graph showing changes in apical glucose with elevations of basolateral glucose ± sucralose ± lactisole; n = 3 ALIs from 3 individual patients used per condition in independent experiments. **(E)** Experiment similar to *(D)* but using only sucralose in ALIs grown from Wt, TAS1R3^-/-^, or GNAT3^-/-^ knockout mice. The ability of sucralose to reduce ASL glucose was lost in TAS1R3^-/-^ mice; n = 3 ALIs per condition in independent experiments. All data are mean ± SEM. Significances in *B*, C, and *D* by one way ANOVA, Bonferroni posttest; \**p*<0.05.

We tested the role of T1R3 in this observation using mouse nasal septal cells grown at ALI from Wt or T1R3 knockout (*TAS1R3*^-/-^) mice.^37^ Sucralose decreased ASL glucose in Wt but not *TAS1R3*^-/-^ mice in the presence of 15 mM basolateral glucose (**Figure 2E**). Because we had shown that T1R3 functions in solitary chemosensory cells, which utilize Gα-gustducin (GNAT3), we tested this response in cells from Gα-gustducin knockout mice (*GNAT3*^-/-^) mice and saw no loss of this sucralose effect (**Figure 2E**), suggesting it is Gα-gustducin-independent.

We hypothesized that a function of T1Rs in airway cilia might be to detect bacterial D-amino acids. Certain D-stereoisomer amino acids have been identified as “sweet” in psychophysics studies^38,39^ and demonstrated to activate the T1R2/3 heterodimer in molecular experiments.^22,32^ It is important to note here that, while we are using the term “sweet,” responses here are distinct from the gustatory sensation of sweetness. We are using this term to refer to activation of the T1R2/3 heterodimer and/or T1R3 homodimer receptor, not the psychophysical perception of taste. To confirm our prior observation^18^ that D-amino acids are present in conditioned media (CM) from patient-derived bacteria composed predominately of *Staphylococcus*, we tested a fluorometric assay that can detect D-amino acids but not their L-isomers (**Figure S2**). We found that CM from clinical and lab (strain M2^40^) *S. aureus* and clinical strains of coagulase-negative *Staphylococcus* (likely *S. epidermidis*) had D-amino acids present. Media from clinical and lab (PAO-1) *P. aeruginosa* strains did not (**Figure 3A**). The D-amino acid content we saw in *Staphylococcus* CM (>1 mM) is similar to the value of ∼1.5 mM reported for *S. aureus* media.^41^ When we looked at CM from sinonasal clinical cultures, we saw D-amino acids present in cultures in which *S. aureus* was predominant but not CM from cultures in which *P. aeruginosa* was predominant (**Figure 3B**). Because these samples came from human patients, they were not strictly single-species cultures, they were classified based on the predominant microorganisms present in each culture, as previously described.^18^ Other species were also likely present in each culture at lower abundance.

**Figure 3.**
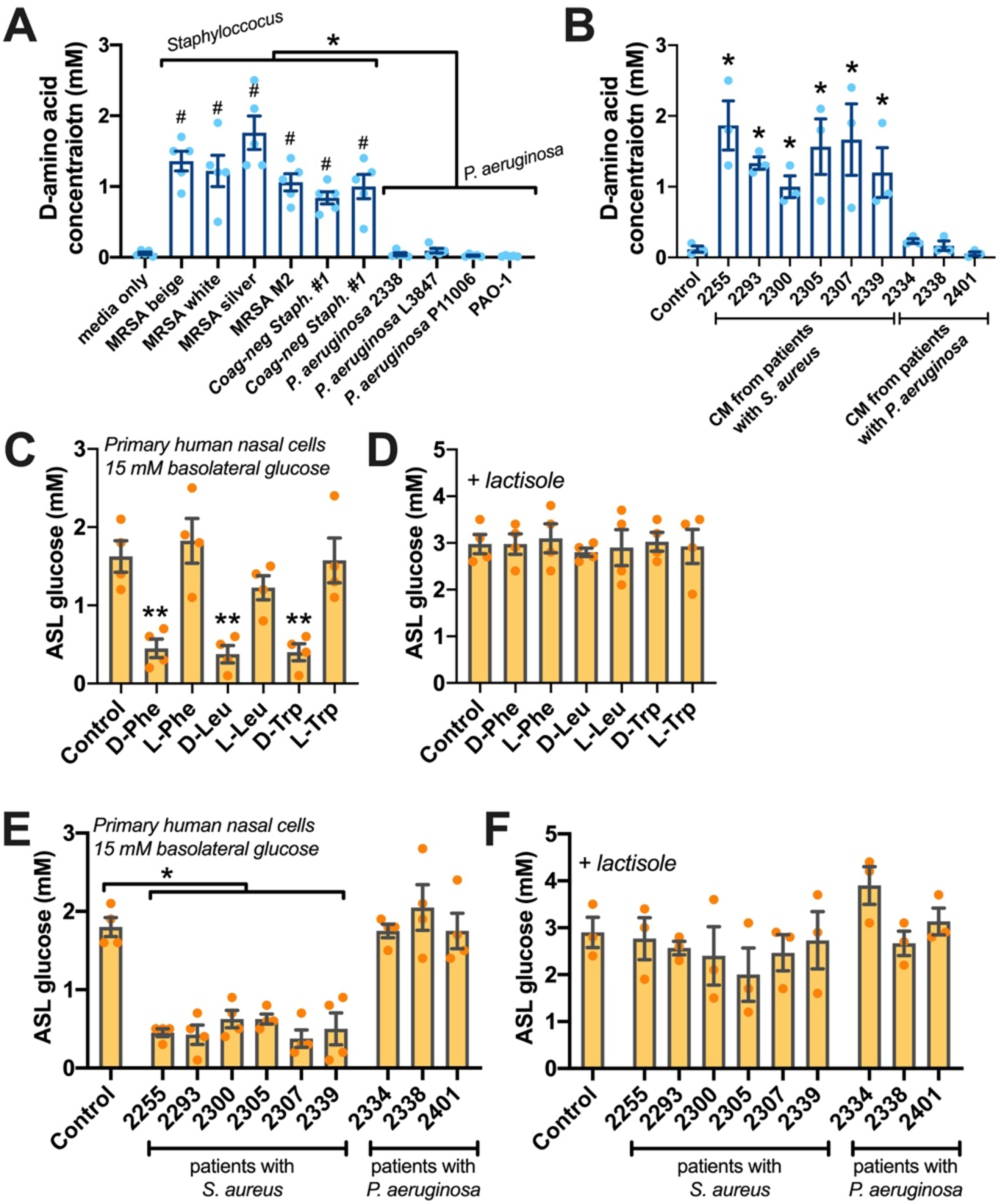
D-amino acids produced by nasal *Staphylococcus* bacteria affect ASL glucose levels in a T1R3-dependent manner. **(A)** Clinical CRS-derived methicillin-resistant *S. aureus* (MRSA; strains named beige, white, or silver) or lab strain MRSA M2 had increased D-amino acid content in their conditioned media. *S. epidermidis* and methicillin-sensitive *S. aureus* likewise produced D-amino acids. No significant D-amino acid production was observed with clinical *P. aeruginosa* strains 2338, L3847, or P11006 or lab strain PAO1. Significance by one-way ANOVA with Tukey Kramer posttest; \**p*<0.05 for *Staphylococcus* vs *P. aeruginosa*; #*p*<0.01 vs control; n = 5 independent experiments per bacteria strain. **(B)** Conditioned media (CM) from patient microbiological cultures predominately containing *S. aureus* produced D-amino acids while those from patients with *P. aeruginosa* did not; n = 3 independent experiments. Significance by one-way ANOVA with Dunnett’s posttest; \**p*<0.05 vs control (LB only). All bar graphs are mean ± SEM. **(C-D)** ASL glucose was measured in 15 mM basolateral glucose used to elevate ASL glucose into a measurable range where reductions could be reliably detected. ASL glucose was reduced by D-Phe, D-Leu, or D-Trp (*A*) but not by L-stereoisomers (*A*) or in the presence of T1R3 inhibitor lactisole (*B*). **(E-F)** ASL glucose (15 mM basolateral glucose) was reduced by CM from patients with *S. aureus* (*E*) but not by CM from patients with *P. aeruginosa* (*E*) or in the presence of lactisole (*F*). All significance determined by one-way ANOVA with Dunnett’s posttest comparing all values to control; \*\**p*<0.01; n = 4 independent experiments using ALIs from 4 individual patients per condition. All bar graphs show mean ± SEM.

Sweet D-amino acids D-Phe, D-Leu, and D-Trp reduced ASL glucose in primary nasal ALIs in the presence of 15 mM basolateral glucose (**Figure 3C**). This effect was reduced in the presence of lactisole (**Figure 3D**). We also saw lowering of ASL glucose with CM from patients infected with *S. aureus* but not with CM from patients with *P. aeruginosa* (**Figure 3E**). Again, this did not occur in the presence of lactisole (**Figure 3F**). Together, these data suggest *S. aureus* D-amino acids might reduce ASL glucose by interacting with T1R3.

### T1R3 enhances apical glucose uptake

To measure glucose uptake, we utilized a fluorescent derivative of D-glucose, 2-[N-(7-nitrobenz-2-oxa-1, 3-diazol-4-yl)amino]-2-deoxy-d-glucose (NBDG). NBDG is transported by most diffusional (GLUT) and sodium linked (SGLT) glucose transporters studied, albeit possibly with slightly altered kinetics.^42,43^ We used NBDG (200 µM) on the apical side only of cultures to preferentially measure cellular uptake from the apical side of the epithelial barrier. We quantified uptake over 70 min of exposure to NBDG (**Figure 4A**) after 18 hours of exposure to apical PBS (control) or PBS with lactisole or sucralose (2 mM each). Prior apical sucralose increased NBDG uptake while lactisole reduced it (**Figure 4A-B**). Acute addition of GLUT inhibitor phloretin (200 µM) or 15 mM glucose to outcompete the NBDG reduced NBDG uptake (**Figure 4A-B**). Use of Na^+^-free (0-Na^+^) PBS had no effect, suggesting NBDG transport is not Na^+^-linked (e.g., SGLT-mediated). GLUT1-specific inhibitor STF-1 (100 µM) had no effect, suggesting the phloretin inhibition was not through GLUT1 (**Figure 4B**). We also saw an increase in NBDG uptake after 18 hours exposure to D-amino acids in the absence but not presence of lactisole (**Figure 4C-D**). L-amino acids had no affect (**Figure 4C**). Prior exposure to CM from patients with *S. aureus* also increased NBDG uptake compared with media only control (**Figure 4E**), and this was reduced in the presence of lactisole (**Figure 4D**). CM from *P. aeruginosa* patients had no effect on NBDG uptake (**Figure 4E**).

**Figure 4.**
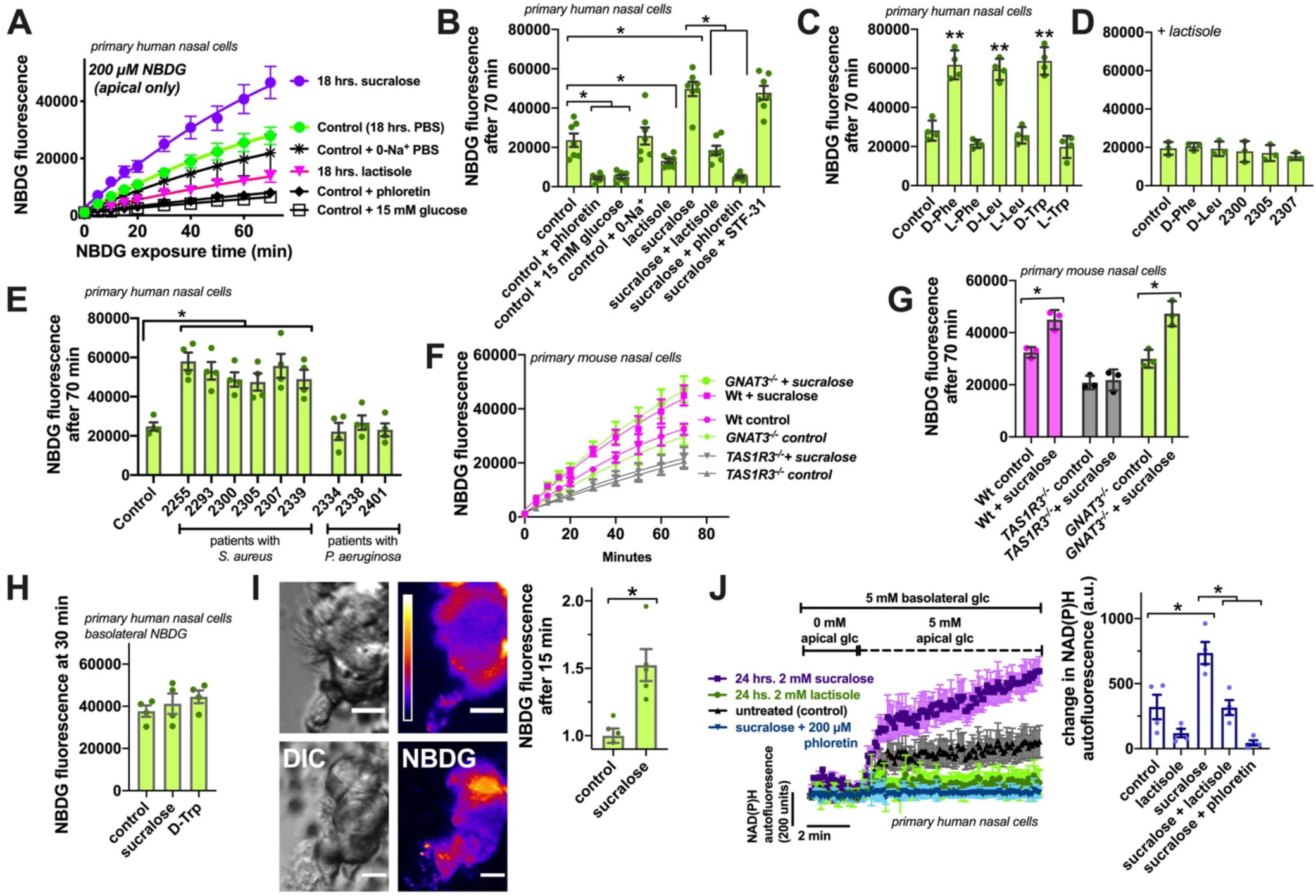
T1R3 regulates apical glucose uptake in nasal epithelial cells. **(A)** Traces showing uptake of fluorescent NBDG (200 µM; apical side only) in cultures pretreated with sucralose or lactisole or in the acute (beginning of the assay) presence of GLUT inhibitor phloretin, 0-Na+ PBS, or 15 mM glucose to outcompete the NBDG. **(B)** Quantification of NBDG fluorescence at 70 min from experiments as shown in *A* (n = 7 individual experiments using ALIs derived from 7 patients). Significance determined by one-way ANOVA with Tukey-Kramer posttest; \**p*<0.05. **(C)** Quantification of apical NBDG uptake at 70 min in the presence of D-isomer or L-isomer amino acids; Significance by one-way ANOVA with Dunnett’s posttest comparing all values to control; \*\**p*<0.01; n = 4 independent experiments using ALIs from 4 different patients. **(D)** Lactisole reduced the increase in apical NBDG uptake stimulated with D-Phe, D-Leu, or D-Trp. Significance by one-way ANOVA with Dunnett’s posttest comparing all values to control + lactisole (all not significantly different); n = 4 independent experiments using ALIs from 4 different patients. **(E)** Bar graph showing increased NBDG fluorescence after 70 min in ALIs exposed to CM from patients with *S. aureus* but not patients with *P. aeruginosa*. Significance by one-way ANOVA with Dunnett’s posttest comparing all values to control; n = 4 independent experiments using ALIs from 4 different patients; \*\**p*<0.01. **(F)** Traces of NBDG fluorescence changes in ALIs derived from Wt, TAS1R3-/-, or GNAT3-/- mice ± sucralose. **(G)** Quantification of NBDG fluorescence at 70 min from independent experiments (n = 3) as in *F*. Sucralose increased apical NBDG uptake in Wt and *GNAT3^-/-^* mice but not *TAS1R3^-/-^*mice. Significance by one-way ANOVA with Bonferroni posttest with paired comparisons; \*\**p*<0.01. **(H)** Bar graph showing lack of difference of basolateral NBDG uptake when applied basolaterally (n = 4 independent experiments using primary nasal ALIs from 4 separate patients). No significant difference by one-way ANOVA. **(I)** Representative images of NBDG fluorescence uptake in cells acutely isolated from brushings of TRPM5-GFP mice, which labels solitary chemosensory cells (SCCs). Cell clumps were visually confirmed to lack SCCs prior to the experiment. Bar graph shows relative change in NBDG uptake after 15 min is increased with sucralose, suggesting this is SCC independent. Significance by Student’s *t* test; \**p*<0.05. **(J)** Left shows representative trace of changes in NAD(P)H autofluorescence during addition of 5 mM apical glucose (in PBS) in cells treated with sucralose or lactisole ± acutely applied (at start of assay) GLUT inhibitor phloretin. Bar graph on right shows quantification of increased autofluorescence changes with sucralose and reduction by lactisole or phloretin. Significance by one-way ANOVA with Tukey-Kramer posttest; \**p*<0.05.

To confirm a role for T1R3, we again utilized cells from *TAS1R3*^-/-^ knockout mice, grown at ALI. There was no increase in NBDG uptake with prior (18 hrs) sucralose treatment in *TAS1R3*^-/-^ ALIs (**Figure 4F-G**). In contrast, both Wt and *GNAT3*^-/-^ ALIs exhibited a similar increase in NBDG uptake after sucralose pre-treatment (**Figure 4F-G**). When NBDG was placed on the basolateral side of cultures, there was no increase in NBDG uptake with sucralose or sweet amino acid D-Trp (**Figure 4H**). Thus, the effects described above are specific for the apical side of the epithelium.

To further test if this is an effect that occurs within ciliated cells or if this requires SCCs, we took cells acutely isolated from mouse nasal turbinate expressing GFP under a TRPM5 promotor,^44^ which labels SCCs (**Figure S3**) as well as taste cells.^13,45^ We used lack of-GFP fluorescence to confirm the absence of SCCs in epithelial clumps. We observed a sucralose-mediated NBDG uptake increase in these cell clumps (**Figure 4I**), suggesting the responses we observed are independent of SCCs. Lactisole was not used here as it does not bind to or inhibit mouse T1R3.^33^

Finally, to confirm the results above with a more physiological assay, we imaged NAD(P)H autofluorescence upon addition of apical glucose after 24 hrs exposure to sucralose ± lactisole ± phloretin. NADH and NADPH, but not NAD and NADP, are autofluorescent when excited by UV light, and addition of metabolites to cells can sometimes be visualized by increases in NAD(P)H production and increase in autofluorescence.^46,47^ Our goal here was not to perform a rigorous metabolic characterization, but rather to test if changes in metabolism could be observed along with the changes in NBDG transport already described. We observed that NAD(P)H autofluorescence increased faster in response to introduction of 5 mM apical glucose cultures pretreated with sucralose and slower in cultures pre-treated with lactisole compared with control cultures (**Figure 4J**). Acute application of phloretin reduced the autofluorescence increase seen with sucralose (**Figure 4J**). These data support the NBDG data suggesting nasal epithelial cells are taking up glucose faster across the apical membrane after pretreatment with sucralose in a lactisole-sensitive fashion. This supports a role for T1R3 in ASL glucose regulation by modifying apical glucose uptake.

### T1R3 increases expression of GLUT2 and GLUT10, likely via arrestin signaling

The major GLUT transporters reported for airway epithelial cells are GLUT2 and GLUT10 on the apical side and GLUT1 on the basolateral side.^2,7^ While one study reported apical GLUT4 in airway epithelial cells,^48^ we did not see substantial GLUT4 transcript expression in nasal ALIs compared with other GLUTs (**Figure 5A**). Publicly available gene expression data also suggest GLUT1 and GLUT10 are the predominant GLUT isoforms in bronchial and nasal cells, with no significant differences in expression with mucociliary differentiation, cigarette smoking, or cystic fibrosis (CF) (**Figure S4**). Thus, we focused largely on GLUT1, GLUT2, and GLUT10 for these studies. Human GLUT10 is a high affinity glucose transporter with KM of ∼0.3 mM,^49–51^ fitting with a potential role in keeping nasal ASL glucose (normally ∼0.5 mM or ∼10-fold below normal serum levels) as low as possible.

**Figure 5.**
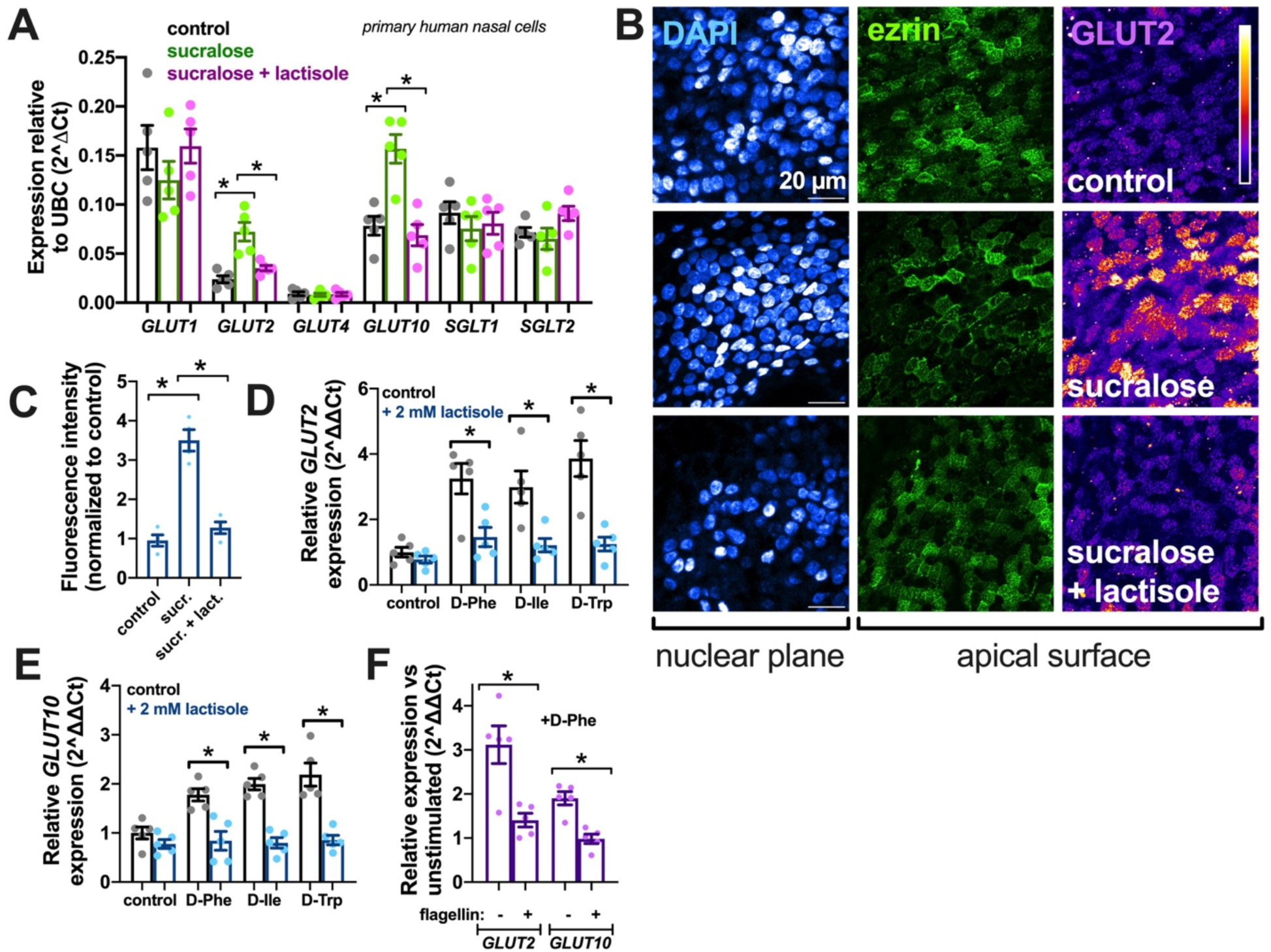
Bacterial D-amino acids increase GLUT2 and GLUT10 expression in a T1R3-dependent manner. **(A)** Expression of GLUT and SGLT isoform transcripts in cells after stimulation with T1R3 agonist sucralose ± T1R3 inhibitor lactisole in primary nasal ALIs. Significance by one-way ANOVA with Bonferroni posttest comparing expression levels of each gene to control conditions; n = 5 independent experiments using ALIs from 5 separate patients per condition; *p*<0.05. **(B)** Images of ALIs after stimulation with sucralose ± lactisole, showing increased GLUT2 immunofluorescence at the apical membrane (indicated by microvillar marker ezrin). **(C)** Bar graph showing quantification of GLUT2 intensity from 4 independent experiments as in *B* using ALIs from 4 individual patients. Significance by one-way ANOVA with Bonferroni posttest; \*\**p*<0.01. **(D-E)** Quantification of GLUT2 (*D*) and GLUT10 (*E*) expression (relative to control conditions) after stimulation with sweet D-amino acids ± lactisole. Significance by one-way ANOVA with Bonferroni posttest with pairwise comparisons; \**p*<0.05. **(F)** Relative expression of GLUT2 and GLUT10 after treatment with D-Phe ± TLR5 agonist flagellin. Significance by one-way ANOVA with Bonferroni posttest with pairwise comparisons; \**p*<0.05.

Sucralose (2 mM; apical side only; 18 hrs;) increased transcript levels of GLUT2 and GLUT10, while GLUT1 transcript was unaltered (**Figure 5A**). The sucralose-induced change in GLUT2 and GLUT10 was blocked by lactisole (2 mM; apical side only; **Figure 5A**). We also observed that sucralose increased GLUT2 immunofluorescence at the apical membrane, identified by microvilli marker ezrin (**Figure 5B-C**). This was also blocked by lactisole (**Figure 5B-C**). Like sucralose, sweet D-amino acids D-Phe, D-Ile, or D-Trp (2 mM; apical side only; 18 hrs), increased GLUT2 (**Figure 5D**) and GLUT10 (**Figure 5E**) in a lactisole-dependent manner. The effects of D-Phe were reduced in cultures pre-treated with flagellin (**Figure 5F**), which reduces T1R3 expression. These data suggest the apical glucose uptake observed above are partly due to GLUT2 and GLUT10 increase.

How does this occur? We previously showed no effect of sucralose on Ca^2+^ signaling, NO production, or ciliary beat frequency in nasal epithelial cells.^13,52^ We again confirmed that there were no detectible changes in these parameters with sucralose or D-Phe treatment in human nasal ALIs (**Figure S5**). We hypothesized the changes in gene expression may occur through β-arrestin-dependent or independent kinase signaling. GLUT2 expression levels may be regulated in pancreatic β cells by β arrestins.^53^ While kinase signaling downstream of taste receptors is grossly understudied, sucralose was previously shown to signal through ERK1/2 signaling in Min6 cells.^54^ We examined primary nasal ALIs after 1 hour of sucralose, D-Phe, or D-Ala stimulation. Cells were fixed and stained with a pan-arrestin antibody. We visualized more intense arrestin localization in cilia with stimulation of sucralose, D-Phe, or D-Ala which was blocked by lactisole (**Figure 6A-B**). Arrestin inhibitor barbadin^55^ reduced GLUT2 and GLUT10 expression increases in response to 18 hrs. sucralose, D-Phe, or D-Ala (**Figure 6C**). No inhibition was observed with phospholipase C inhibitor U73122 (1 µM), Gαi inhibitor pertussis toxin (10 ng/ml), or rho-kinase inhibitor Y-27632 (1 µM) (**Figure 6D**), suggesting cilia T1R3 responses are independent of G protein signaling and perhaps tied to arrestin-mediated kinase signaling. Supporting this, when cultures were treated with siRNAs against β-arrestins 1 and 2, we found that the D-Phe-stimulated increase in GLUT2 and GLUT10 expression was blocked (**Figure 6E**).

**Figure 6.**
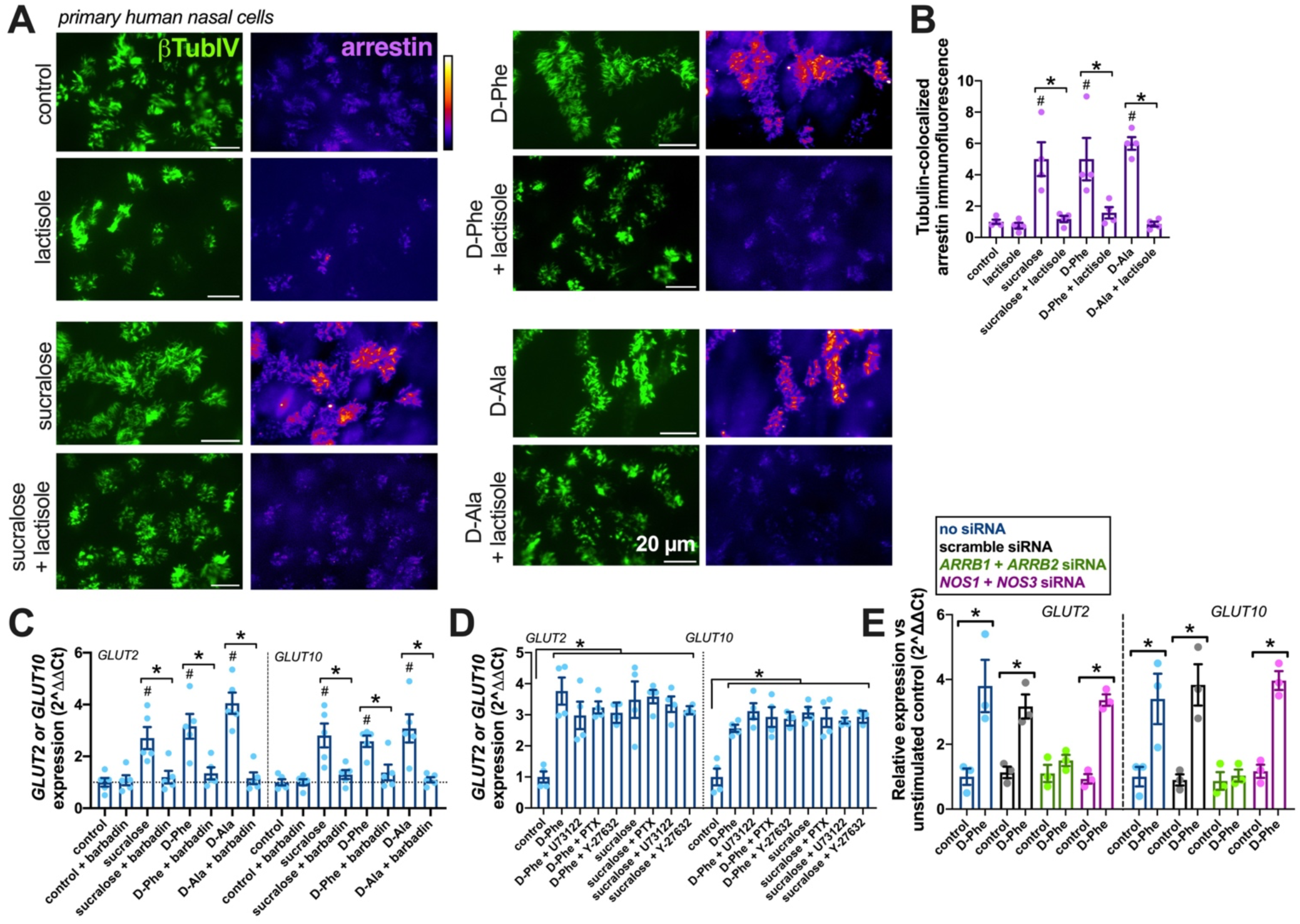
T1R3 regulation of GLUT2 and GLUT10 expression likely involves arrestin signaling. **(A)** Immunofluorescence images of primary nasal cilia after stimulation with sweet agonists sucralose, D-Phe, or D-Ala ± lactisole. Cilia marker (β-tubulin IV) and pan-β-arrestin primary antibodies were used. All sets of images taken at identical microscope settings. **(B)** Quantification of cilia-localized (β-tubulin IV-co-localized) arrestin immunofluorescence from 4 independent experiments as in *A* using cells from 4 separate patients. Significance by one-way ANOVA with Bonferonni posttest comparing all values to control as well as comparing each condition ± lactisole; ^#^*p*<0.05 vs control; \**p*<0.05 between bracketed bars. **(C)** *GLUT2* and *GLUT10* expression in cultures stimulated with sucralose, D-Phe, or D-Ala ± β-arrestin/β2-adaptin inhibitor barbadin. Significance by one-way ANOVA with Bonferonni posttest comparing all values to control as well as comparing each condition ± barbadin; ^#^*p*<0.05 vs control; \**p*<0.05 between bracketed bars. **(D)** *GLUT2* and *GLUT10* expression in cultures stimulated with sucralose or D-Phe ± PLC inhibitor phospholipase C inhibitor U73122, Gαi/Gαgustducin inhibitor pertussis toxin (PTX), or rho kinase inhibitor Y-27632. Significance determined by one-way ANOVA with Bonferroni posttest comparing all values to control (\**p*<0.05) or suraclose or D-Phe only as appropriate. Only the control group was p<0.05 vs sucralose or D-Phe only. **(E)** Experiments were carried out to measure *GLUT2* or *GLUT10* expression in human nasal ALI cultures after treatment with no siRNA (blue), scramble siRNA (black), siRNA against β-arrestins 1 and 2 (ARRB1 and ARRB2; green), or endothelial and neuronal NOS isoforms (NOS 1 and NOS 2; magenta). Cultures were then unstimulated (control) or stimulated as in C-D with D-Phe. Significance between control vs D-Phe determined by one-way ANOVA with Bonferroni posttest with paired comparisons; *p<0.05.

### ASL glucose protects *S. aureus* against epithelial NO innate defenses

While others have shown that *S. aureus* can use glucose as a carbon source,^4^ we confirmed that addition of 0.5 mM glucose can enhance bacterial growth even in a nutrient rich media (DMEM + 1× MEM amino acids; **Figure S6**). Thus, the mechanism described above may serve an innate immune role. Namely, increasing glucose mucosal uptake may serve to limit bacterial growth by denying them an ASL nutrient source. However, we were also intrigued by a study showing that glycolysis can confer bacterial resistance to antimicrobial NO production.^56^ *S. aureus* is known to be able to detoxify NO and resist this innate defense mechanism.^57^ We previously found that *S. aureus* are more resistant to NO than gram-negative *P. aeruginosa*.^58^ Additionally, nasal cilia T2Rs detect bitter bacterial lactones and quinolones from gram-negative bacteria to activate nitric oxide (NO) production to clear and kill bacteria.^27–29^ However, others have shown that gram positive bacteria can also activate T2Rs.^59^ We previously showed that *S. aureus* conditioned media (CM) activates nasal epithelial NO production, though this was not tied to a specific receptor.^60^

Using the fluorescent reactive nitrogen species indicator DAF-FM, we confirmed that there was increased RNS production upon stimulation of human nasal ALIs with MRSA M2 conditioned media (**Figure 7A**). This likely reflected NO production as it was blocked by pre-treatment (10 µM; 45 min) with NOS inhibitor L-NAME but not inactive D-NAME (**Figure 7A**). This likely represents activation of an innate defense NO pathway in response to metabolites or other components produced by the *S. aureus*, possibly mediated by T2R activation. We also observed increased DAF-FM fluorescence with CM from clinical MRSA isolates (**Figure 7B**).

**Figure 7.**
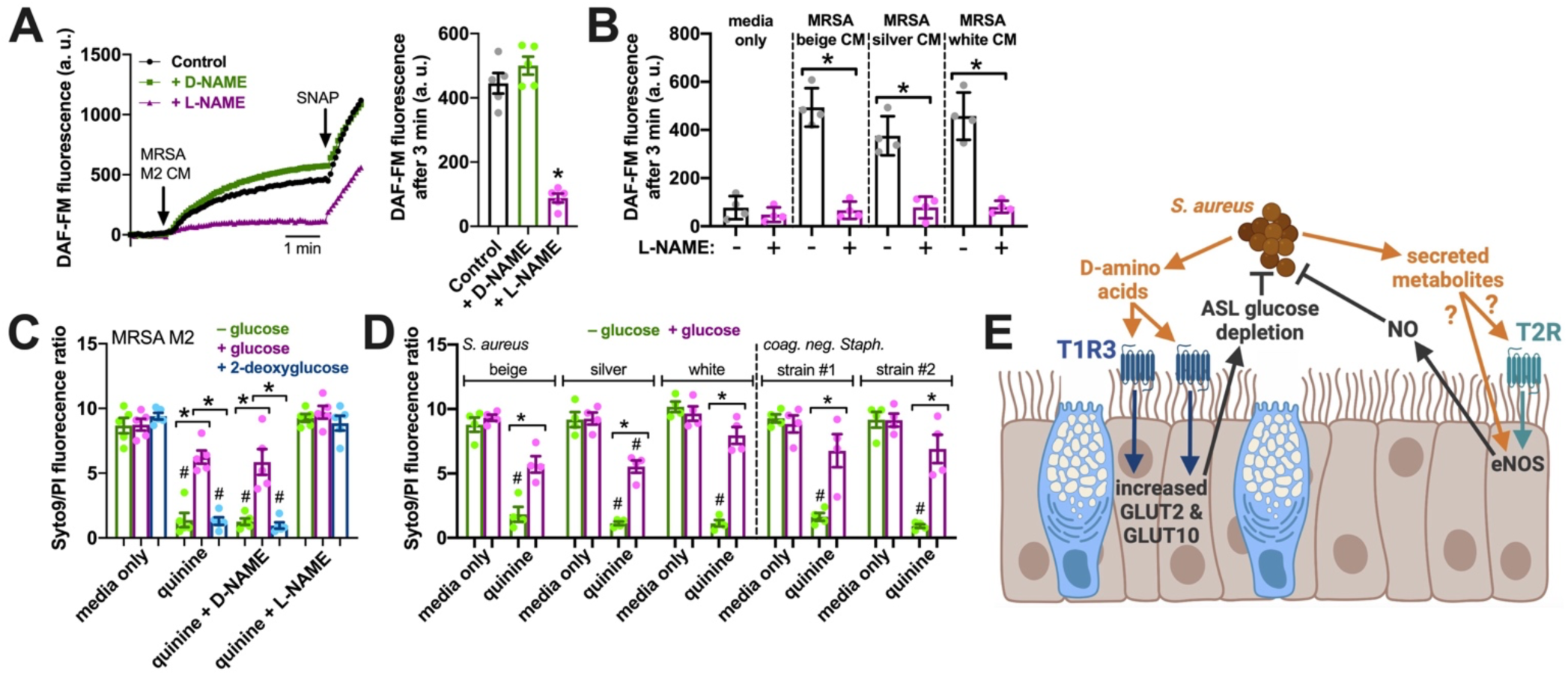
Glucose protects *S. aureus* against T2R-dependent NO innate defense. **(A)** Representative DAF-FM traces (left) and bar graph (right; n = 5 independent experiments using ALIs from 5 patients) showing NO production during MRSA M2 conditioned media (CM) exposure. As a control, cultures were pretreated with NOS inhibitor L-NAME or inactive D-NAME (10 µM, 45 min). Significance vs control (no L-NAME or D-NAME) determined by one-way ANOVA with Dunnett’s posttest; *p<0.05. **(B)** Bar graph of DAF-FM fluorescence after 3 min stimulation with CM from clinical MRSA isolates ± NOS inhibitor L-NAME. Significance between bracketed bars determined by one-way ANOVA with Bonferroni posttest for paired comparisons; \**p*<0.05. **(C)** Bar graph of the ratio of SYTO9 (live) vs propidium iodide (PI; dead) fluorescent stain emission from MRSA M2 bacteria in co-culture bacterial killing assays as described in the text. Quinine was used at 0.1 mg/ml, which alone had no effect on bacteria. Green bars show results from cultures with no apical glucose, magenta bars show results from cultures with 0.5 mM apical glucose, and blue bars show results from cultures with 0.5 mM 2-deoxyglucose. **(D)** Results from another set of experiments similar to C but in media ± quinine ± glucose and using clinical S. aureus strains or two strains of coagulase-negative *Staphylococcus* (likely *S. epidermidis*). Significance in *C* and *D* determined by one-way ANOVA with Bonferroni posttest with pre-defined comparisons; *p<0.05 vs bracketed group and ^#^p<0.05 vs media only. Results from n = 5 independent experiments using a total of 4-5 cultures from 4-5 different patients per condition. **(E)** Working model of cilia-T1R3 regulation of innate defense in response to *S. aureus* D-amino acids. We hypothesize that D-amino acids are detected by T1R3 in airway cilia, which increases GLUT2 and GLUT10 expression to increase ASL glucose uptake and reduce ASL glucose concentration. This ASL glucose depletion likely limits pathogen growth. However, ASL glucose depletion also enhances the sensitivity of *S. aureus* to bactericidal NO produced downstream of T2R bitter receptors or other pathogen recognition mechanisms. Image created with Biorender.

We hypothesized that elevated ASL glucose may protect *S. aureus* from the bactericidal effects of NO production downstream of T2R activation by allowing the bacteria to use glycolysis to detoxify the NO, as described.^56,61^ When MRSA M2 were washed and resuspended in glucose-free DMEM media and incubated with ALI cultures in a previously developed NO-dependent bactericidal assay,^27,52,62^ we found that addition of T2R agonist quinine^63^ stimulated bacterial killing over 2 hours, evidenced by increased propidium iodide (dead cell stain) fluorescence compared with Syto9 (live cell stain) fluorescence (**Figure 7C**). This was NO dependent as it was blocked by L-NAME but not D-NAME (**Figure 7C**). The addition of 0.5 mM glucose reduced bacterial killing (**Figure 7C**). However, equimolar 2-deoxyglucose, which cannot undergo glycolysis, did not reduce bacterial killing (**Figure 7C**). Similarly, when we incubated clinical *S. aureus* strains as well as coagulase-negative *Staphyloccocus*, we found that the presence of glucose also inhibited quinine/NO-activated bacterial killing (**Figure 7D**). Together, these data suggest that the presence of metabolizable glucose in the ASL protects *Staphyloccocus* from bactericidal NO produced during T2R stimulation in nasal cells

## DISCUSSION

Here we show that T1R3 taste receptor subunit is expressed in nasal epithelial cilia and detects bacterial “sweet” D-amino acids and regulates expression of glucose transporters GLUT2 and GLUT10, likely through an arrestin-dependent pathway. This is a new role for T1R3 in the airway epithelium. It suggests that activation of T1R3 by *S. aureus* D-amino acids enhances glucose uptake to reduce ASL glucose and reduce bacterial growth (**Figure 7E**, left). Reduced ASL glucose may also enhance *S. aureus* NO-mediated defenses activated by T2Rs or other detection mechanisms (**Figure 7E**, right).

On the tongue, the sweet taste receptor is likely a heterodimer of T1R2 and T1R3.^64–66^ It remains to be determined if T1R2 and/or umami receptor subunit T1R1 plays a role in the airway cilia response observed here. T1R3 homodimers have been suggested to function in pancreatic beta cells,^67–69^ which could also be the case here. Future work is needed to determine the dimerization partner(s) of T1R3 in cilia, though this is partially limited by the availability of reliable T1R antibodies. The use of T1R3 knockout mice and T1R3 antagonist lactisole in this study was critical to rigorously determining a role for T1R3. While we previously detected T1R1, T1R2, and T1R3 in differentiated nasal epithelial cell ALIs by qPCR, expression of T1R2 was at least a log lower than T1R1 or T1R3.^70^

While the exact mechanism(s) of the T1R3 signaling to regulate GLUT2/GLUT10 expression changes also remains to be further determined, our data suggest it might be independent of G protein signaling and instead dependent on arrestin-mediated kinase signaling. Further work is needed to explore the relatively unknown non-G-protein arms of taste receptor signaling. One novel aspect of this work is that it suggests that airway motile cilia chemosensation involves more than just the cAMP, Ca^2+^, and NO pathways previously implicated in the signaling from these organelles.^24,25,52^

We hypothesize that a sweet agonist (artificial sweetener), delivered via topical nasal rinse, might reduce ASL glucose via the mechanisms described in our study and be beneficial in the management of CRS and other sinonasal pathology. The identity of potential agonists with efficacy partly depends on whether other T1R isoforms are expressed. While sugars like sucrose and glucose likely bind to the extracellular venus fly trap domains of both T1R2 and T1R3,^71–73^ some artificial sweeteners like aspartame^32,74,75^ and xylitol^76^ bind to T1R2 and others like cyclamate bind to T1R3.^31^ Future screening is needed to determine the most efficacious sweet agonists, which could be characterized using the differentiated human nasal ALI model used here. Differences exist between human and mouse sweet taste receptors that might preclude use of mice as a screening or preclinical model. Mice have reduced ability to detect some artificial sweeteners due to sequence differences between human and mouse T1R2.^74,77^ Likewise, differences in the T1R3 sequence cause antagonist lactisole to not bind to mouse T1R3.^33^ While our data suggest mechanistic studies might be performed in mice, drug screening studies should likely employ human cells.

It also remains to be determined if this glucose regulation system is truly a *bona fide* global defensive response to limit pathogen growth or a more subtle way to change the ASL to a more supportive environment for some bacteria and less supportive environment for others. Glucose is an excellent carbon sources for many bacteria, including *Staphylococcus* gram-positive species and gram-negatives like *Escherichia coli* and *P. aeruginosa*.^78,79^ However, *E. coli* and *P. aeruginosa* may grow better in high glucose conditions compared to *Staphylococcus aureus* and commensal *Staphylococcus epidermidis*.^80^ The downregulation of this T1R3 cilia pathway by *P. aeruginosa* flagellin-TLR5 signaling (reducing T1R3 ASL glucose regulation) and converse activation of the T1R3 cilia pathway by *S. aureus* D-amino acids suggests that different bacteria can differentially modulate this pathway. Whether this is host-pathogen or host-commensal/host-microbiome crosstalk remains to be elucidated. It might be both, as determined by the level of the relative stimuli the host cell receives.

However, the fact that glucose metabolism can affect *S. aureus* susceptibility to NO suggests the implications of lowering ASL glucose reach further than simply limiting nutrients for bacteria growth. NO is likely an ancient and conserved mechanism of host-pathogen interactions at ciliated epithelia. It was recently shown that NO is a key component of the recruitment of bioluminescent *Vibrio fischeri* to the developing bobtail squid light organ, lined by ciliated cells that produce NO in response to the bacteria.^81^ In the airway, limiting glucose may specifically be beneficial in *S. aureus* infections by enhancing their ability to be killed by NO. To our knowledge, the intersection of nasal ASL glucose and effectiveness of NO bacterial killing is a novel innate immune crosstalk pathway in the airway. This may be specifically relevant to diseases like CF, where airway NO production is reduced and ASL glucose can be elevated with the onset of CF-related diabetes.^62^

One question that arises from our study is whether delivery of a topical sweet agonists to activate cilia T1R3 might be detrimental by subsequent inhibition of SCCs. This may indeed be the case and warrants further investigation. However, SCCs do not appear to be as common in the human airway as in mice, at least in healthy airways. However, detailed rigorous studies of SCC presence and localization in the human airways are still to be undertaken. SCCs appear to be regulated by type II cytokines like IL-13 or IL-4 and/or exposure to fungal metabolites.^16,82^ Thus, SCCs may only be prevalent in the nose of CRS patients with nasal polyps (CRSwNP), a subset of CRS driven by type II cytokines,^83^ or in patients with allergic fungal sinusitis. CRS without (*sans*) nasal polyps (CRSsNP) is most often characterized by other inflammatory profiles.^83^ We might find that sweet agonists are useful in CRSsNP but not CRSwNP. Alternatively, activating T1Rs to suppress SCCs in CRSwNP may also suppress SCC IL-25 generation and reduce inflammation and still be beneficial despite suppression of antimicrobial production. Determining how cilia T1R3 fits into the context of these diseases will require several more *in vivo*-focused studies. However, differential TAS1R3 single nucleotide polymorphism (SNP) distributions were observed in control, CRSsNP, and CRSwNP populations,^84^ hinting that this receptor may be important in upper respiratory diseases.

In summary, our data reveal T1R3 as a novel regulator of ASL glucose that could be targeted in CRS or other airway diseases. We show a new interkingdom signaling pathway where bacterial D-amino acids are detected by T1R3 to regulate glucose transport. This pathway likely cross-talks with nasal epithelial NO production to indirectly control the bactericidal effects of nasal NO through bacterial metabolism.

## Acknowledgements

The authors thank M. Victoria, L.E. Kuek, J.R. Freund, and B. Chen (University of Pennsylvania) as well as G. Xiong (Children’s Hospital of Philadelphia) for technical assistance and helpful discussions. We thank Dr. L. Chandler (Philadelphia VA Medical Center) for microbiology assistance and isolation of clinical strains. This study was supported by NIH/NIDCD grant R01DC016309 to R.J.L., NIH/NIAID grant R01AI167971 to N.D.A., J.N.P, and R.J.L., and a pilot project grant to R.J.L. from the Penn Diabetes Research Center (P30DK19525).

## CRediT authorship contribution statement

Ryan M. Carey: Resources, Conceptualization, Data curation, Investigation, Methodology, Writing – original draft, Writing – review & editing. Benjamin M. Hariri, Investigation, Methodology; Nithin D. Adappa: Resources, Data curation, Project administration, Writing – review & editing. James N. Palmer: Resources, Data curation, Project administration, Writing – review & editing. Robert F. Margolskee: Resources, Methodology, Writing – review & editing. Noam A. Cohen: Resources, Conceptualization, Data curation, Project administration, Writing – review & editing. Robert J. Lee: Conceptualization, Investigation, Methodology, Formal analysis, Writing – original draft, Writing – review & editing, Project administration, Funding aquisition

## Data Availability

All data are contained within the manuscript. All materials will be provided upon request to the corresponding author.

## STAR ★ METHODS

### KEY RESOURCES TABLE

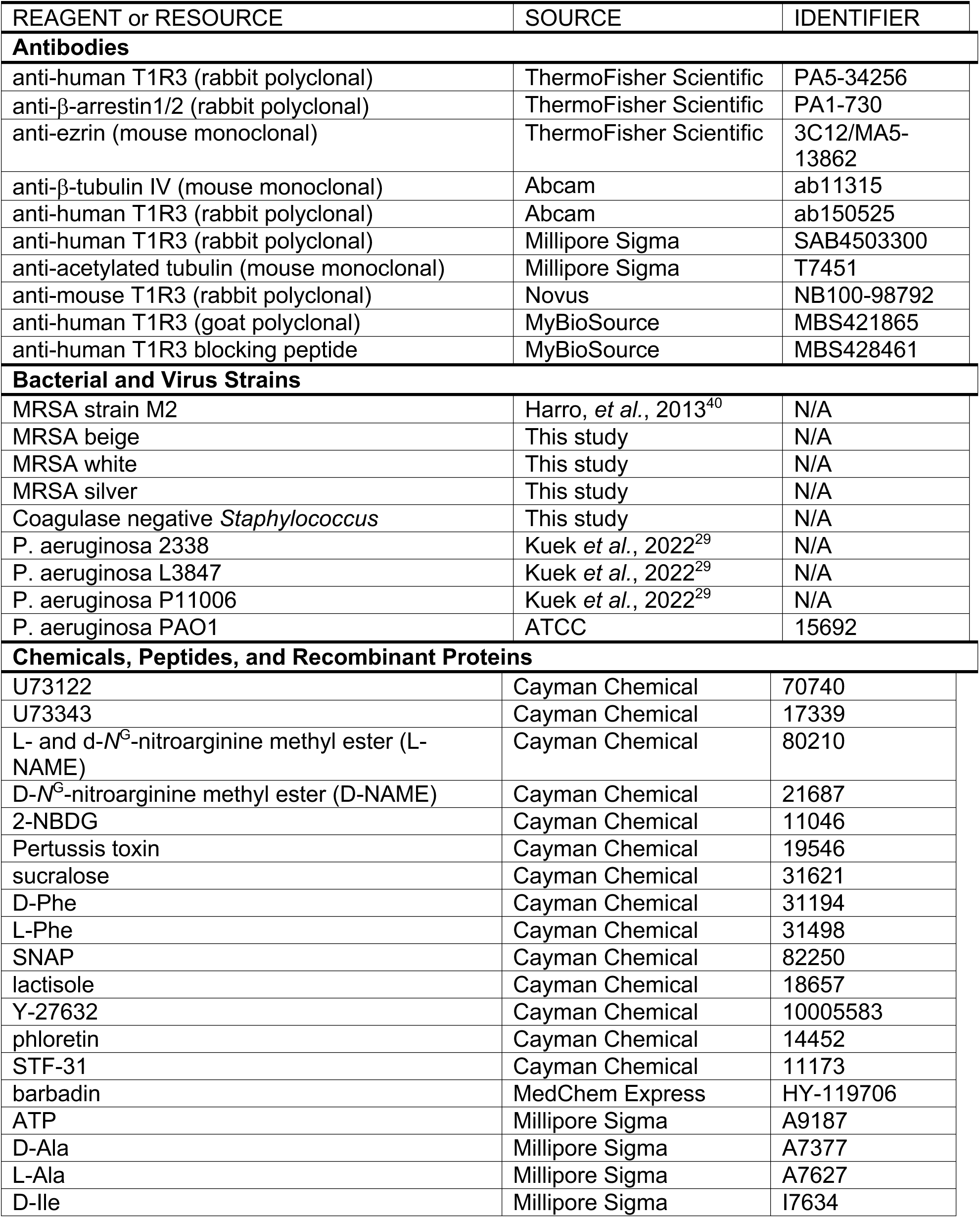
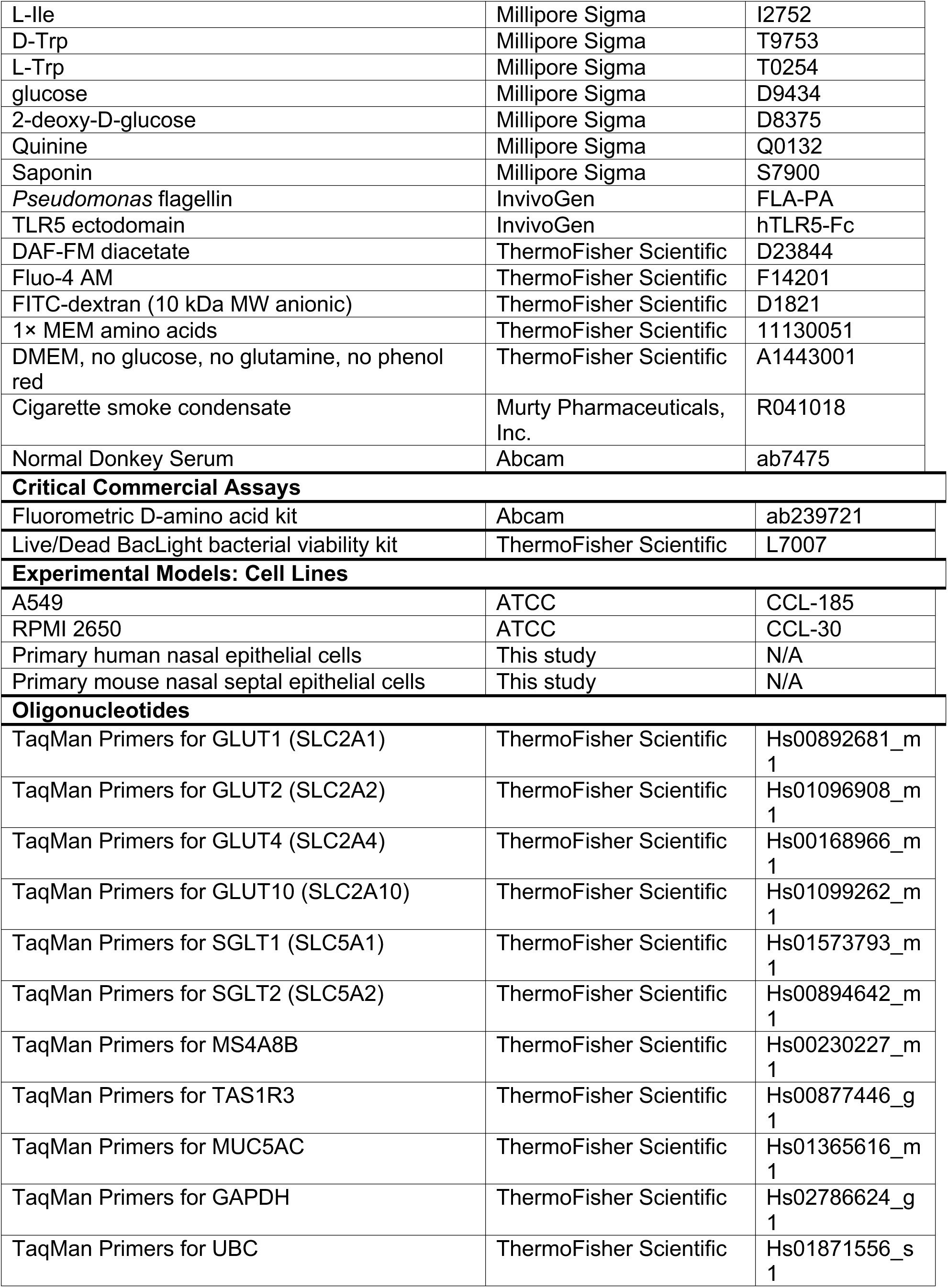
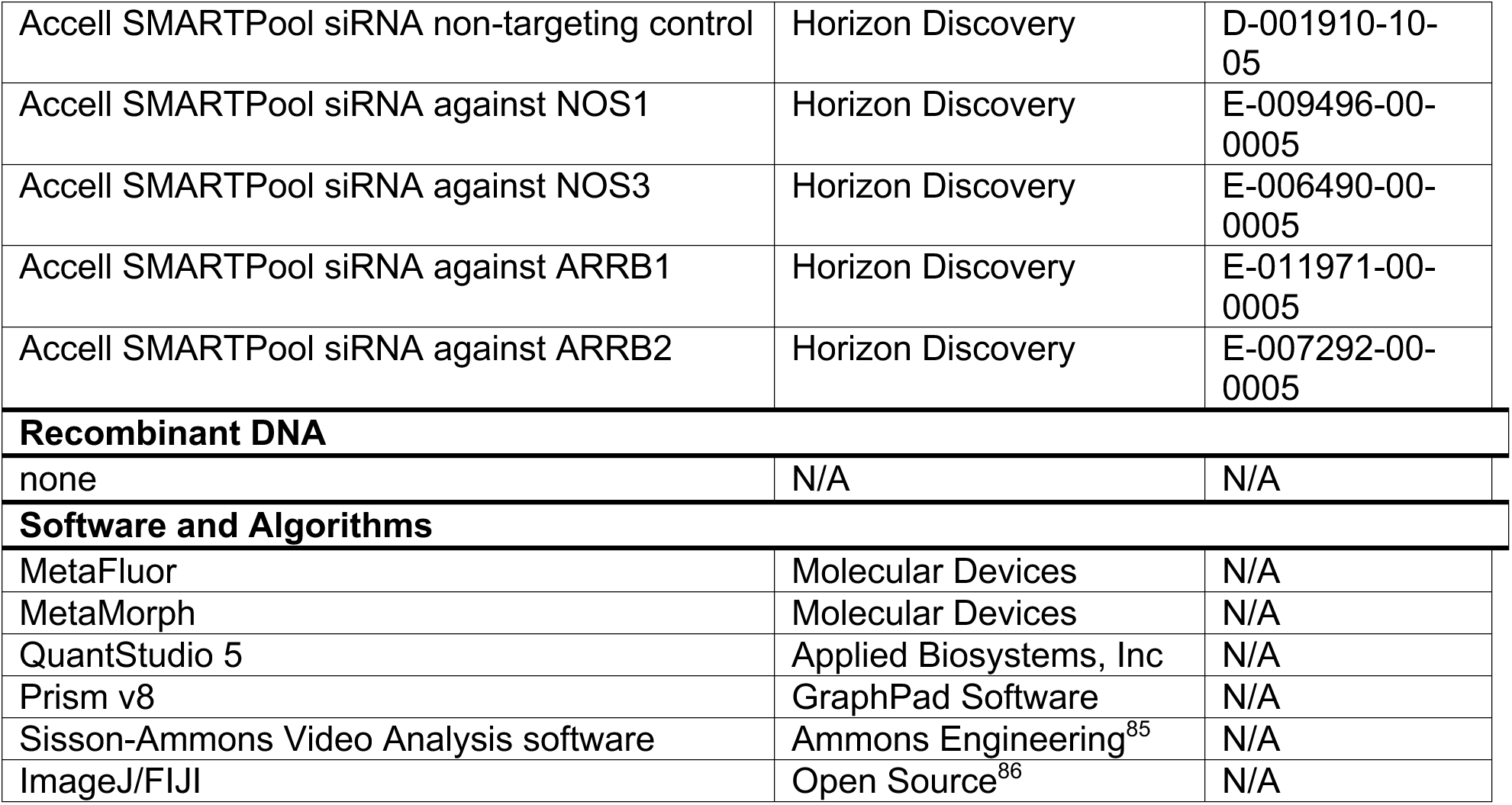

### RESOURCE AVAILABILITY

#### Lead contact

Robert J. Lee.

#### Materials availability

Further information and requests for resources and reagents should be directed to and will be fulfilled by Robert J. Lee.

#### Data and Code availability

The published article includes all datasets generated or analyzed during this study. Raw numerical values used to generate bar graphs or traces are available upon request.

### EXPERIMENTAL MODEL AND SUBJECT DETAILS

#### Cell culture

Immortalized cells (RPMI2650 and A549) were obtained directly from ATCC and grown in minimal essential media (MEM; Gibco/ThermoFisher Scientific) plus 1× cell culture penicillin/streptomycin and 10% FetalPlex serum substitute (Gemini Biosciences). For air-liquid interface (ALI) cultures, cells were seeded onto collagen-coated 0.33 cm^2^ transwell filters (Corning) and grown to confluence for 5 days before apical air exposure, as described previously ^87,88^. Cultured cells were used after 10-14 days at ALI. TEER was measured using an epithelial volt-ohm meter (EVOM; World Precision Instruments) and chopstick electrodes. Routine mycoplasma testing of all cell lines was carried out by the University of Pennsylvania Cell Center Service Facility.

Primary human nasal surgical specimens were obtained according to The University of Pennsylvania guidelines for use of residual clinical material from patients undergoing sinonasal surgery at the Hospital of the University of Pennsylvania with institutional review board approval (#800614). Written informed consent was obtained from patients in accordance with the U.S. Department of Health and Human Services code of federal regulation Title 45 CFR 46.116. Patients ≥18 years of age requiring surgery for sinonasal disease or trans-nasal approaches to the skull base were included. Patients with systemic inheritable disease (eg, granulomatosis with polyangiitis, systemic immunodeficiencies) or those using antibiotics, oral corticosteroids, or anti-biologics (e.g. Xolair) within one month of surgery were excluded. Vulnerable populations (patients ≤18 years of age, pregnant women, and cognitively impaired persons) were also excluded.

Mucosal tissue was transported to the lab on ice in saline for subsequent enzymatic dissociation. Cells were grown to confluence in proliferation medium (7 days), then lifted and seeded at high density on transwells (Corning transparent, 0.33 cm^2^, 0.4 µm pore size) as described.^52,88–90^ The next day, basolateral media was changed to differentiation medium and culture medium was removed from the upper compartment. Differentiation media contained 1:1 Lonza bronchial epithelial cell basal media (BEBM):Dulbecco’s modified Eagle’s medium (DMEM) plus Lonza singlequot supplements (0.5 ng/ml hEGF, 5 ng/ml epinephrine, 0.13 mg/ml BPE, 0.5 ng/ml hydrocortisone, 5 ng/ml insulin, 6.5 ng/ml triiodothyronine, and 0.5 ng/ml transferrin, 0.1 nM retinoic acid) supplemented with 1× penicillin/streptomycin and 2% NuSerum (BD Biosciences/Corning, San Jose, CA) as described.^52^ Cells were fed from the basolateral side only with differentiation media for ∼21-28 days before use.

Mouse nasal septal ALI cultures were grown as described^91^ from residual tissue from animals sacrificed for other experimental purposes with IACUC approval. No mice were sacrificed for the purposes of this study. Epithelial cells were isolated by collagenase and pronase digestion before culture on Costar 6.5 mm transwell permeable filter supports (Corning Inc. Life Sciences, Lowell, MA USA) with apical side submerged. After 7 days, cells reached confluence and the medium was removed from the apical surface with feeding from the basolateral side. Differentiation and ciliogenesis were observed within 10–14 days after exposure to air, and cultures were used within 4–6 weeks.

### METHOD DETAILS

#### Solutions

Dulbecco’s phosphate buffered saline (DPBS) contained (in mM) 138 NaCl, 2.7 KCl, 1.5 KH2PO4, 8 Na2HPO4, 1.8 CaCl2, and 1.5 MgCl2, with pH adjusted to approximately 7.2. Hank’s balanced salt solution (HBSS) used for live cell imaging and other experiments contained (in mM): 138 NaCl, 5.3 KCl, 0.34 Na2HPO4, 0.44 KH2PO4, 0.49 MgCl2, 0.41 MgSO4, 1.3 CaCl2, 20 HEPES free acid. HBSS was adjusted to pH 7.4 with NaOH. Glucose (5.5 mM) was added unless indicated otherwise. When 1x MEM amino acids was added to the HBSS, pH was adjusted after addition of the amino acids. For all assays, basolateral media was changed to phenol-free glucose-free Dulbecco’s modified Eagle’s medium (DMEM; Gibco/ThermoFisher Scientific) plus 2% Corning NuSerum and glucose added to the indicated basolateral concentrations.

#### Immunofluorescence microscopy

Immunofluorescence staining for cilia was carried out as described.^52,88,90^ ALI cultures fixed in 4% formaldehyde for 20 min at room temperature, followed by blocking and permeabilization for 1 hour at 4°C in Dulbecco’s phosphate buffered saline (DPBS) containing 1% bovine serum albumin (BSA) plus 5% normal donkey serum (NDS) to block non-specific protein binding and 0.2% saponin plus 0.3% Triton X-100 to permeabilize. After three washes (5 min each) in DPBS, primary antibody incubation (1:100) was performed in DPBS containing BSA, NDS, and saponin overnight at 4°C. After another round of washing in DPBS, incubation with AlexaFluor (AF)-conjugated donkey anti-mouse or anti-rabbit secondary antibodies (1:1000; ThermoFisher Scientific) was performed at 4°C for 2 hours in DPBS plus BSA, NDS, and saponin.

After staining, transwell filters were removed from plastic retaining rings and mounted onto slides with Fluoroshield + DAPI mounting media (Abcam; Cambridge, MA USA). For co-staining with direct labeling of primary antibodie, Zenon antibody labeling kits (Thermo Fisher Scientific) for AF546 or AF647 were used as described.^52,88,90^ In some experiments, images of ALIs were taken on an Olympus Fluoview confocal system with IX-73 microscope and 60× (1.4 NA) objective. For other experiments, images were taken on an Olympus IX-83 microscope with spinning disk confocal unit (Olympus DSU) with 60× (1.4 NA) objective using Metamorph. All images were analyzed in FIJI ^86^ using only linear adjustments (min and max), set equally between compared images obtained at identical microscope settings (exposure, objective, binning, etc.).

#### ASL glucose measurements

ASL glucose concentrations were measured as previously described.^13^ ALI cultures were transferred to 5.5 mM glucose (minimal essential media + Earl’s salts + 1% NuSerum) for 24 hrs. Basolateral glucose was then changed to the values indicated in the text, and 30 μl PBS was placed on the apical side of the ALIs for 24 hours, assuming that glucose concentration would equilibrate over this time and reflect the physiological ASL concentrations generated by these cultures. Samples were collected and run on a colormetric glucose assay (Cayman Chemical) using manufacturer-provided glucose standards.

#### NBD-glucose uptake

2-NBDG in PBS was incubated on the apical side of cultures at the concentrations and times indicated in the text. Followed by incubation, cells were copiously washed with NBDG-free PBS and immediately imaged in the center of the culture using standard FITC filter set on an Olympus IX83 microscope with 4× 0.16NA objective and Metamorph (Molecular Devices). Images were taken at exactly the same microscope settings with no nonlinear adjustments.

#### NAD(P)H autofluorescence

NAD(P)H autofluorescence^46,47^ was imaged as described.^92^ We excited ALIs with a Xenon lamp (Sutter Lamda LS 300 W) with high UV output at 340 nm and measured fluorescence at 450 nm using a DAPI filter set and 30× 1.0 NA silicone oil immersion objective lens with high UV transmittance on an IX-83 microscope (Olympus) equipped with a Hammamatsu Orca Flash 4.0 sCMOS camera (Hammamatsu, Tokyo, Japan). Images were acquired every 12 seconds using Metafluor (Molecular Devices, Sunnyvale CA). Transwells with no cells were used to subtract background from plastic autofluorescence. Cultures were kept in 5 mM basolateral glucose, and apical surface contained 30 µL of 0-glucose PBS. Apical glucose was changed from 0 mM to 5 mM by subsequent addition of an equal volume of PBS supplemented with 10 mM glucose.

#### Quantitative reverse-transcription PCR (qPCR)

Quantitative reverse transcription PCR was carried out as described.^29,93^ ALIs were lysed in TRIzol (ThermoFisher) and RNA was isolated using Direct-zol (Zymo Research). cDNA was transcribed via High-Capacity cDNA Reverse Transcription Kit (ThermoFisher) and gene expression was quantified utilizing Taqman qPCR probes and QuantStudio 5 Real-Time PCR System (ThermoFisher). Data were analyzed using Excel and GraphPad PRISM. UBC was used as a housekeeping control gene as described.^29,93^

#### Primary nasal ALI siRNA experiments

For siRNA, primary nasal ALIs were treated with Acell SMARTPool siRNAs against indicated genes or non-targeting pool (Catalog ID: D-001910-10-05) as per the manufacturer’s instructions using a protocol previously described^94^ and utilized by our lab.^62^

#### Measurement of FITC-dextran flux

FITC-dextran protocol was based on^95^ and previously carried out in our lab.^96^ Briefly, FITC-dextran (10 kDa) in phenol-red-free glucose-free DMEM was placed on the apical side of the culture with sucralose ± lactisole, and basolateral solution (phenol red-free glucose-containing DMEM) was collected from the basolateral side after 30 min of incubation at 37 °C. Samples were immediately read in 96 well fluorescence plates using a Tecan Spark 10M fluorescence plate reader (485 nm excitation, 525 nm emission). Basolateral media from cells treated with apical media only (containing no FITC-dextran) were used for background measurement and subtraction.

#### Microbiological cultures

Patient sinonasal microbiology cultures were collected from sinonasal swabs using BBL CultureSwab Plus transport system (Becton, Dickinson, and Co., Sparks MD USA), grown overnight in lysogeny broth (LB) media for 24 hrs at 37 °C, followed by freezing of glycerol stocks and speciation by the Philadelphia VA Medical Center microbiological laboratory (Dr. L. Chandler). Conditioned media was obtained from subsequent 24 hr overnight cultures normalized by diluting turbidity to 0.5 MacFarland and filtering through a 0.2 µm filter. Sweet D-amino acid production was previously demonstrated from the *Staphylococcus* patient CMs used here via liquid chromatography and tandem mass spectrometry (LC-MS/MS).^18^ This was confirmed by D-amino acid assay used here.

For preparation of conditioned medium (CM), *S. aureus* strains were grown for 24 hours at 37°C with shaking in LB. The 24-hour overnight culture was than diluted to 0.1 optical density (OD) (log phase), and grown under the same conditions for 12 hours. The resultant medium was centrifuged at 2000g for 10 minutes at room temperature and filtered through a 0.2-μm filter. CM was used at a final concentration of 25%, diluted in Dulbecco’s phosphate-buffered saline (DPBS); this concentration was used because the equivalent concentration of LB medium was the highest concentration that did not evoke Ca^2+^ or NO responses itself (this study and references^52,60,97^).

For anti-bacterial assays (conducted as described^27,52,62^), bacteria were grown to OD 0.1 in LB and resuspended in glucose-free, pyruvate-free DMEM media with or without 0.5 mM glucose. Previously saline was used in this assay but DMEM was used to give bacteria some nutrients in the absence of glucose. Data in Figure S6 show minimal effects on bacterial growth over 2 hrs., which is the duration of the assay. This ensures results are not caused by starving the bacterial. Moreover, the Syto9/PI method is ratiometric, and thus is independent of the exact number of bacteria but rather reflects the live-dead ratio or general health of the population. Nasal ALIs were washed 24 hrs prior with antibiotic-free Ham’s F12K media (ThermoFisher Scientific) on the basolateral side. Bacteria suspension (30 uL) was then placed on the apical side of the ALI for 10 min, followed by aspiration of bulk fluid. Following incubation for 2 hrs at 37°C, the remaining bacteria were removed from the ALI washing. This was followed by life-dead staining with SYTO9 (live) and propidium iodide (dead) (BacLight Bacterial Viability Kitl; ThermoFisher Scientific; cat # L7012). Green (live)/red (dead) ratio was quantified in a Spark 10M (Tecan, Mannedorf, Switzerland) at 485 nm excitation and 530 nm and 620 nm emission.

For planktonic growth assays, MRSA strains were cultured in LB (ThermoFisher) at 37 °C with shaking overnight. Bacteria were spun down and washed in glucose-free DMEM media then diluted to OD 0.1 in 10 mL total volume DMEM + 1× MEM amino acids with or without added 1 mM glucose. Cultures were grown at 37 °C with shaking (180 RPM); 1 mL of solution was removed at each time point and assayed for OD at 600 nm in a spectrophotometer.

#### Ca^2+^ and NO imaging

Fluo-4 (Ca^2+^ indicator) and DAF-FM (NO indicator) were imaged as previously described^62,88,90^ using a FITC filter set. Images were aquired using MetaFluor (Molecular Devices, Sunnyvale, CA USA) on an IX-83 microscope (10× 0.4 NA PlanApo objective) with xenon lamp excitation source (Sutter Lambda LS, Sutter Instruments, Novato, CA USA) and Orca Flash 4.0 sCMOS camera (Hamamatsu, Tokyo, Japan). All experiments utilized background measurements of unloaded ALIs taken at the same microscope settings; background measurements were subtracted from experimental fluorescence values from each individual wavelength recorded for each experiment.

Primary human nasal ALIs were loaded for 2 hrs in the dark with 10 µM DAF-FM-diacetate or fluo-4 acetoxymethyl ester (AM) in 20 mM glucose-free HEPES-buffered HBSS plus 0.1% pluronic F127 on the apical side. The basolateral side of the culture was incubated in HBSS containing glucose supplemented with 1× MEM amino acids on the basolateral side, followed by washing three times with the same buffer. Compounds were added to the apical side in glucose-free HBSS.

#### Imaging and quantification of ciliary beat frequency (CBF)

Whole-field CBF was imaged using a Basler A602 camera and phase-contrast Nikon TS-100 microscope (40x long working distance objective) was imaged as described^62,88,90^ at 120 frames per second at ∼26-28°C in a glass bottom chamber. Dulbecco’s PBS (+1.8 mM Ca^2+^) was used on the apical side and 20 mM HEPES-buffered Hank’s Balanced Salt Solution supplemented with 1× MEM vitamins and amino acids was used on the basolateral side. Data were analyzed using the Sisson-Ammons Video Analysis software^85^ and normalized to baseline CBF.

#### Data analysis and statistics

Numerical data were compiled in Excel (Microsoft) and graphed in Prism (GraphPad software, La Jolla, CA). All data in bar graphs are shown as mean ± SEM. Statistical analyses of multiple comparisons were made in Prism using one-way ANOVA with Bonferroni (pre-selected pairwise comparisons) or Dunnett’s (comparing to control value) post-tests; *p* <0.05 was considered statistically significant, as indicated by asterisks (*) or pound signs (^#^). Raw data points used to construct graphs or traces are available upon request.

