## Supplemental Data for "Taste receptor T1R3 in nasal cilia detects *Staphylococcus aureus* D-amino acids to increase apical glucose uptake and enhance innate immunity"

### SUPPLEMENTAL MATERIAL

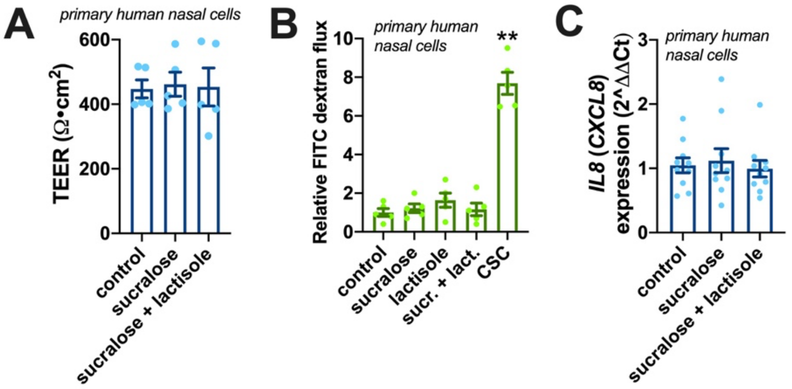


**Figure S1: Lack of effect of sucralose and lactisole on nasal ALI permeability or inflammatory status. (A)** Graph showing no change in TEER with sucralose or lactisole (n = 5 ALIs from 5 different patients per condition in independent experiments). No significant differences by one-way ANOVA. **(B)** FITC dextran permeability showed no change in permeability with sucralose, lactisole, or sucralose + lacitsole. Cigarette smoke condensate (CSC; 40 μg/ml) was used as a positive control for barrier breakdown[96] (n = 5 ALIs from 5 different patients per condition in independent experiments). Only CSC was different compared with control by one-way ANOVA with Dunnett's posttest; **p<0.01. **(C)** Graph showing no change in *IL8/CXCL8* expression by qPCR with sucralose or lactisole (n = 10 ALIs from 10 different patients per condition in independent experiments). No significant differences by one-way ANOVA.

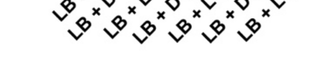


**Figure S2: D-amino acid quantification.** Validation of D-amino acid assay to detect D- but no L- stereoisomer amino acids. Significance by one-way ANOVA with Dunnett's posttest; ***p*<0.01 vs LB only control; n = 3 independent experiments.

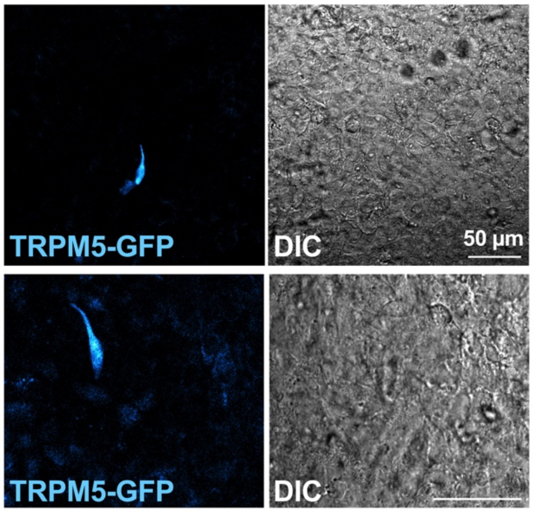


**Figure S3: TRPM5-GFP-labeled mouse nasal ALIs.** Representative laser scanning confocal images from two mouse nasal septal ALIs showing GFP-labeled TRPM5 positive solitary chemosensory cells (SCCs),[44,45,98] also known as tuft cells. We previously showed that human SCCs also express T1R3 and also detect bacterial D-amino acids.[18] Differential interference contrast (DIC) image of the intact epithelium is shown on the right.

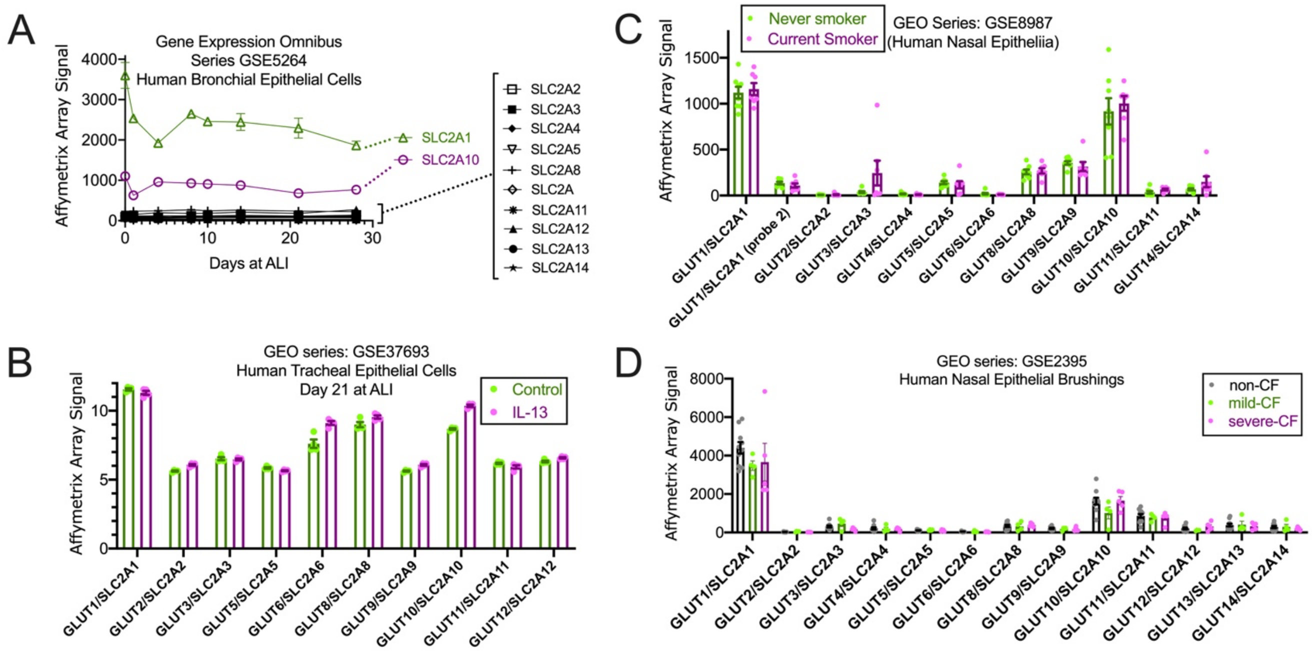


**Figure S4. Gene expression data of GLUT isoforms in bronchial and nasal epithelial cells. (A)** Time course showing GLUT (SLC26A family) expression in human bronchial air-liquid interface (ALI) cultures over 28 days of mucociliary differentiation from Ross, *et al*.[99] SLC2A1 (GLUT1) and SLC2A10 (GLUT10) were the highest GLUTs expressed. **(B)** Comparison of GLUT/SLC2A expression in tracheal ALIs ± IL-13 from Alevy, et al.[100] No significant differences by one-way ANOVA. **(C)** Comparison of GLUT expression from current and never smokers, from Sridhar, et al.[101] No differences by one-way ANOVA. **(D)** Comparison of GLUT expression in non-CF vs CF patients from Wright, et al.[102] No significant differences by one way ANOVA.

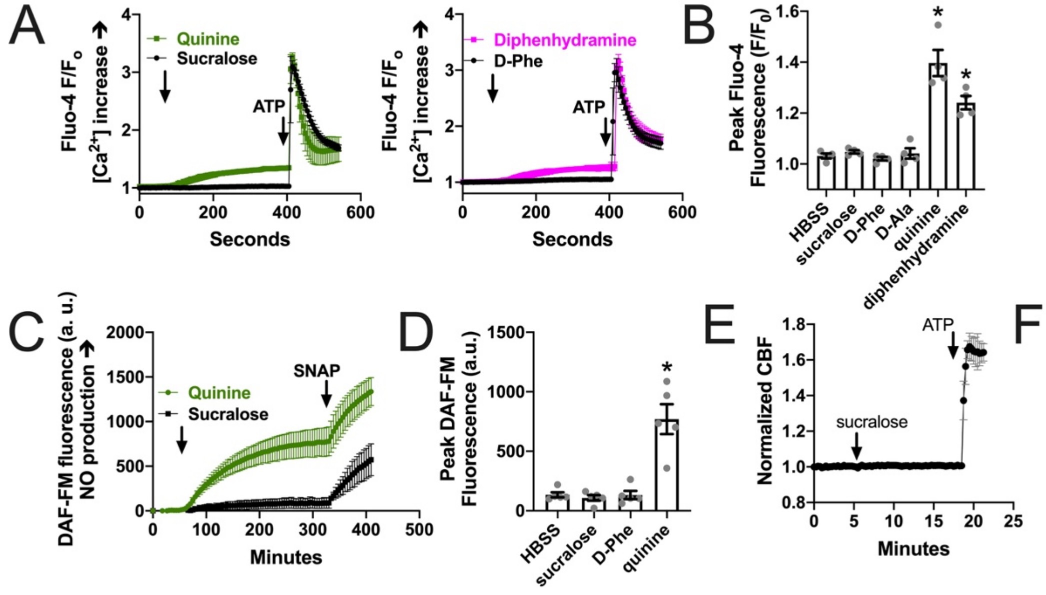


**Figure S5. Lack of effect of sucralose or D-Phe on $Ca^{2+}$ signaling, nitric oxide (NO) signaling, or ciliary beat frequency (CBF). (A)** Representative fluo-4 $Ca^{2+}$ traces during stimulation with T2R agonists quinine (600 μM), diphenhydramine (5 mM), or sucralose (5 mM) or D-Phe (5 mM). Purinergic agonist ATP (100 μM) is used as a positive control at the end of the experiment. **(B)** Bar graph of peak $Ca^{2+}$ with T1R and T2R agonists from A from 4 independent experiments using ALIs from 4 patients. Buffer only (Hank's balanced salt solution, HBSS) was used as a control. Significance vs control determined by one-way ANOVA with Dunnett's posttest; *$p$<0.05. **(C)** Representative DAF-FM traces showing NO production in response to quinine (600 μM) or sucralose (5 mM). NO production with quinine was verified by NOS inhibition (L-NAME vs D-NAME) and NO scavenging (cPTIO) in previous studies.[63,90] **(D)** Bar graph showing results from experiments as in *C* from 5 independent experiments using ALIs from 5 patients. Significance vs HBSS control determined by one-way ANOVA with Dunnett's posttest; *$p$<0.05. **(E)** Representative ciliary beat frequency (CBF) traces showing lack of response to sucralose. Purinergic agonist ATP (100 μM) used as a positive control. **(F)** Bar graph showing results from experiments as in *E* from 5 independent experiments using ALIs from 5 patients. Significance vs HBSS control determined by one-way ANOVA with Dunnett's posttest; *$p$<0.05.

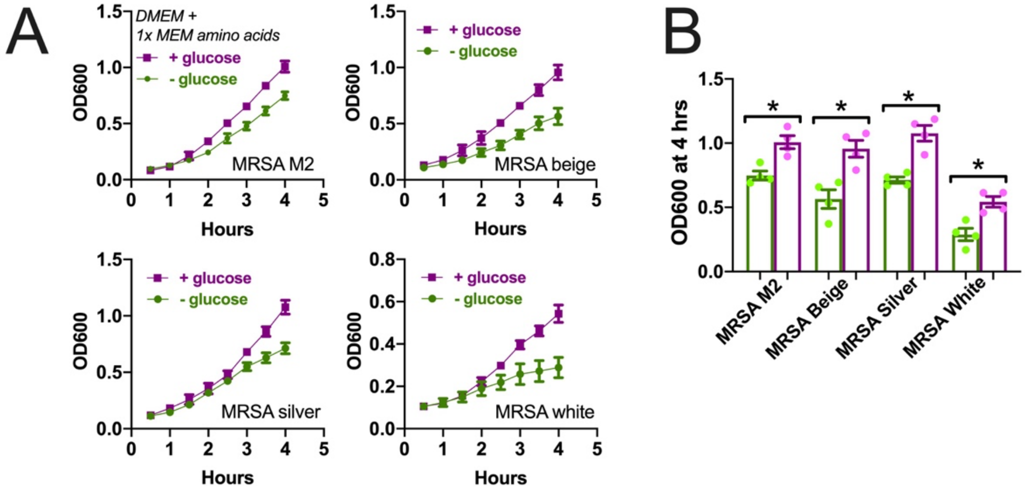


**Figure S6. Glucose enhances the growth of *S. aureus* in nutrient-rich but glucose-free DMEM media.** Methicillin-resistant *S. aureus* strains (lab strain M2 and clinical isolates beige, silver, and white) were cultured as described[29] except using glucose-free phenol-red-free DMEM media supplemented with additional 1× MEM amino acids to create a nutrient rich media lacking glucose to test the effects of 0.5 mM glucose on cell growth. **(A)** Curves showing planktonic growth in glucose (magenta) or no glucose (green) conditions over four hours from a starting OD of 0.1. Shown are mean ± SEM of 4 independent experiments. **(B)** Bar graph of data at 4 hours from A, showing significant increase of growth in the presence of glucose. Significance by one-way ANOVA with Bonferroni posttest for paired comparisons; *$p<0.05$.
